# SARS-CoV-2 Omicron XBB lineage spike structures, conformations, antigenicity, and receptor recognition

**DOI:** 10.1101/2024.02.12.580004

**Authors:** Qianyi E. Zhang, Jared Lindenberger, Ruth J. Parsons, Bhishem Thakur, Rob Parks, Chan Soo Park, Xiao Huang, Salam Sammour, Katarzyna Janowska, Taylor N. Spence, Robert J Edwards, Mitchell Martin, Wilton B. Williams, Sophie Gobeil, David C. Montefiori, Bette Korber, Kevin Saunders, Barton F. Haynes, Rory Henderson, Priyamvada Acharya

## Abstract

A recombinant lineage of the SARS-CoV-2 Omicron variant, named XBB, appeared in late 2022 and evolved descendants that successively swept local and global populations. XBB lineage members were noted for their improved immune evasion and transmissibility. Here, we determine cryo-EM structures of XBB.1.5, XBB.1.16, EG.5 and EG.5.1 spike (S) ectodomains to reveal reinforced 3-RBD-down receptor inaccessible closed states mediated by interprotomer receptor binding domain (RBD) interactions previously observed in BA.1 and BA.2. Improved XBB.1.5 and XBB.1.16 RBD stability compensated for stability loss caused by early Omicron mutations, while the F456L substitution reduced EG.5 RBD stability. S1 subunit mutations had long-range impacts on conformation and epitope presentation in the S2 subunit. Our results reveal continued S protein evolution via simultaneous optimization of multiple parameters including stability, receptor binding and immune evasion, and the dramatic effects of relatively few residue substitutions in altering the S protein conformational landscape.

## Introduction

Since its discovery in November 2021, the SARS-CoV-2 Omicron B.1.1.529 variant, noted for its high number of mutations in the spike (S) protein relative to previous variants of concern (VOC), has continued to mutate and has diversified into many sublineages that have gone through a series of global shifts in dominance (**Figure S1**) (Hadfield et al., 2018). Identified in late 2022, the XBB lineage, a recombinant of two highly diversified BA.2 sublineages, became the first documented example of a SARS-CoV-2 variant increasing its fitness through recombination rather than substitutions. XBB.1, the early XBB lineage member that gave rise to the initial expansion of the XBB lineages, was remarkably resistant to antiviral humoral immunity induced by vaccination or breakthrough infections of prior Omicron subvariants, with its immune resistant properties attributed to multiple receptor binding domain (RBD) substitutions and the Y144del N-terminal domain (NTD) mutation that has been recurrently observed and originally noted in the Alpha VOC (Tamura et al., 2023). XBB.1 was more infectious than BA.2 in pseudovirus infectivity assays and its RBD bound to ACE2 stronger than BA.2 RBD. While significantly more fusogenic than BA.2 and BA.2.75, XBB.1 pathogenicity was comparable to BA.2.75 and less than Delta (Tamura et al., 2023).

XBB.1.5, a descendent of XBB.1 harboring an additional F486P substitution in the RBD, was identified around October 2022. XBB.1.5 became the most prevalent lineage worldwide outcompeting the previously most prevalent Omicron BA.5 descendent BQ.1.1 variant (**Figure S1**) (Uriu et al., 2023). As XBB.1.5 began to decline by mid-summer 2023, XBB.1.16 together with a complex array of additional minor lineages replaced it. Around July 2023, EG.5 began to rapidly spread in some Asian and North American countries (Kaku et al., 2023). EG.5’s higher effective reproduction number (R0) than XBB.1.9.2 (parental with same S protein amino acid sequence as XBB.1.5), XBB.1.5 and XBB.1.16, attributed to its enhanced immune evasion properties, contributed to its global spread. EG.5.1, the immediate descendent of EG.5, had one additional Q52H substitution in the NTD (Wang et al., 2023), and was the EG.5 sublineage that spread most rapidly.

As expected from their sequence similarities, the S proteins of XBB lineage members, XBB.1.5, XBB.1.16 and EG.5, cluster together on a phylogenetic tree and are distinct from other VOCs and from their parental BA.2 (**Figure S2**). Given the lead time required for vaccine evaluation and manufacture, XBB.1.5, the dominant variant in the spring of 2023, was selected as the S variant to be used as a monovalent booster for the vaccine update (Pfizer, Moderna, Novavax) that become available in the fall of 2023. (FDA, 2023a, FDA, 2023b). Early studies have shown the booster to be protective against currently circulating variants (Link-Gelles et al., 2024).

The S protein on the SARS-CoV-2 surface, composed of S1 and S2 subunits, is the primary determinant of its receptor binding and immune evasion properties (**Figure S2**). The S1 subunit is responsible for receptor binding and forms a cap over the fusion subunit S2 in the pre-fusion conformation of the S protein. The S1 NTD subunit (residues 27–305) is connected via a linker (N2R; residues 306–334) to the RBD (residues 335–521), followed by the subdomain 1 (SD1; residues 529–591) and subdomain 2 (SD2; residues 592–697) regions. The RBD contains a receptor binding motif (RBM; residues 483–506) that binds host receptors including ACE2 (Jackson et al., 2022) and TMEM106B (Baggen et al., 2023). The RBD adopts either a receptor-inaccessible “down” or closed conformation, or a receptor-accessible “up” or open conformation. SD2 harbors a furin cleavage site that defines the boundary between the S1 and S2 subunits. Another cleavage site S2’ is located in the S2 subunit. Following receptor binding and proteolytic processing of the S protein, the S2 subunit reorganizes upon detachment of S1, allowing for insertion of the hydrophobic fusion peptide into the host membrane and transition of the S2 helices to the postfusion conformation, leading to cell entry.

The early Omicron variants BA.1 and BA.2 acumulated stabilizing mutations at the RBD-RBD interprotomer contacts in the receptor-inaccessible down state (Gobeil et al., 2022, Stalls et al., 2022, Zhang et al., 2023), resulting in a shift of the S protein population towards greater occupancy of the receptor-inaccesible 3-RBD-down state, either through direct interactions between the RBDs or by stabilizing structural features involved in interprotomer RBD interactions. These mutations are retained in the XBB.1.5, XBB.1.16 and EG.5 S proteins (**Figures 1A and S2**). To understand how the S protein structures changed in these evolved Omicron subvariants, here we determine cryo-EM structures and study the receptor binding, antigenicity and stability of XBB.1.5, and of the closely related XBB.1.16, EG.5 and EG.5.1 S ectodomains using our previously established S-GSAS-D614G (uncleaved) or S-RRAR-D614G (furin-cleaved) platforms prepared without the incorporation of any proline stabilization mutation, as well as a construct expressing only the RBD of each variant (**Figure S2, Data S1 and S2**) (Gobeil et al., 2021a, Gobeil et al., 2022, Stalls et al., 2022, Gobeil et al., 2021b).

**Figure 1.**
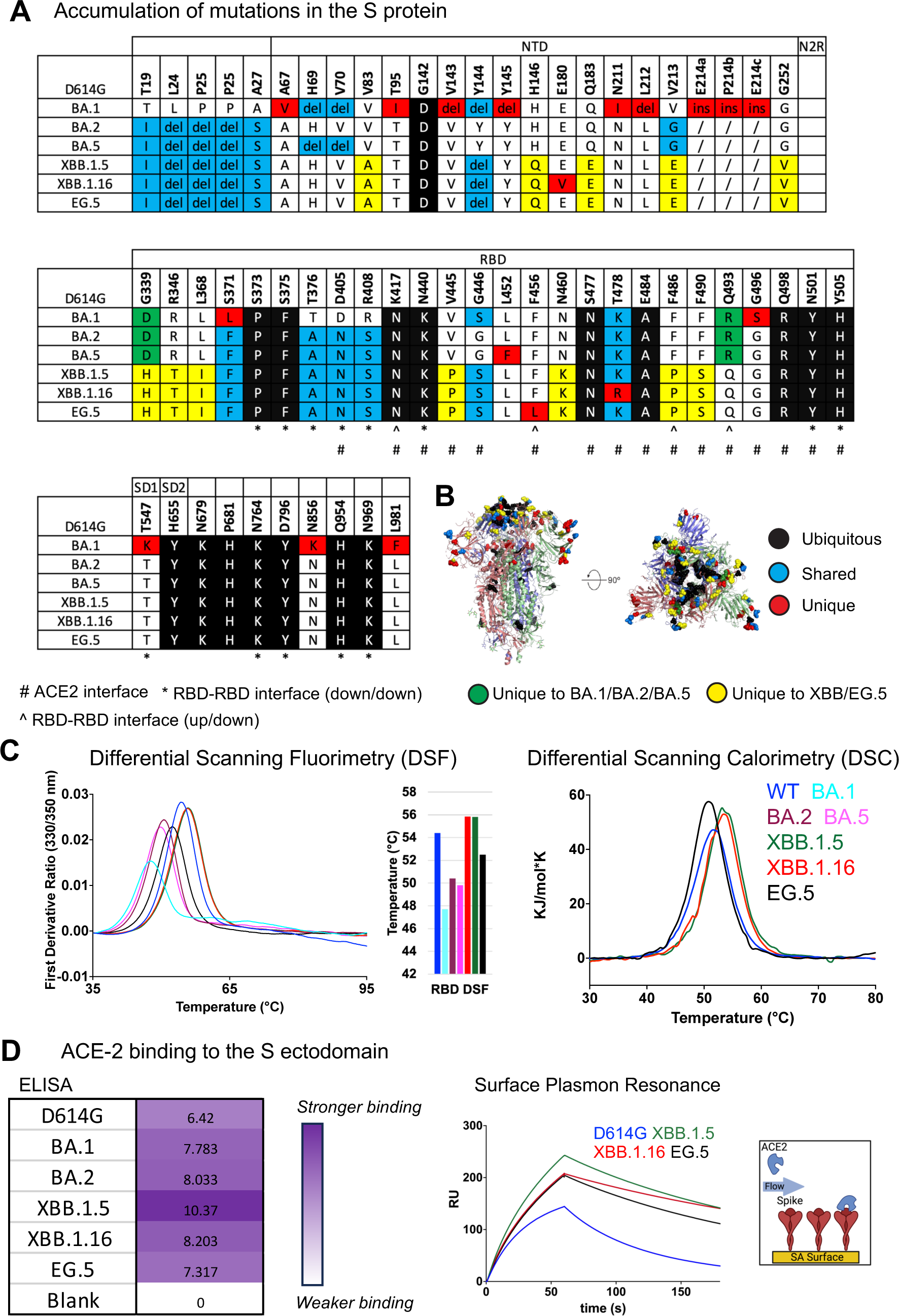
Evolution of SARS-CoV-2 Omicron S protein, RBD stability, and receptor binding. **A.** Mutations in the S protein shown relative to the D614G variant. The first row indicates the positions of the major domains on the S protein. Mutations that are unique to BA.1, BA.2 and BA.5 but do not occur in XBB.1.15, XBB.1.16 and EG.5 are colored green. Those that are unique to XBB.1.5, XBB.1.16 and EG.5 but do not occur in BA.1, BA.2 or BA.5 are colored yellow. Mutations that occur in all the variants listed are termed “Ubiquitous” and colored black. Those that occur in at least two of the variants are called “Shared” and colored cyan, those that are occur in only one variant are called “Unique” and colored red. The del and ins indicate deletion and insertion, respectively. **B.** Positions of mutations shown in panel A marked on a BA.2 S protein structure (PDB 7UB0). The three S protein protomers are colored salmon, blue and light green. The mutations are shown as spheres with the same color scheme as in panel A. **C.** Left. DSF profiles of RBD showing changes in protein intrinsic fluorescence (expressed as the first derivative of the ratio between fluorescence at 350 and 330 nm) with temperature. A bar graph shows the inflection temperature for each RBD variant plotted along the Y-axis. Right. DSC profiles of RBD showing changes in heat capacity with temperature. **D.** Binding of S ectodomain to the ACE2 receptor ectodomain. Left. ELISA Area under the Curve (AUC) shown for each variant and color coded in different shaded of purple according to a scale that indicates strong binding (dark purple) to no binding (white). Right. ACE2 binding measured using Surface Plasmon Resonance (SPR). The S protein constructs used encoded 1-1208 residues with the furin site mutated to a GSAS sequence.

## Results

### Thermostability of the XBB.1.5, XBB.1.16 and EG.5 RBD

New mutations in the XBB.1.5, XBB.1.16, EG.5 and EG.5.1 S proteins relative to the original Omicron lineages BA.1, BA.2 and BA.5 are located in the NTD and RBD, with several appearing at the ACE2 interacting site (**Figures 1A** and **B**). We have previously reported dramatic loss in BA.1 RBD stability relative to the WT RBD, with subsequent BA.2 RBD mutations partially recovering RBD stability (Stalls et al., 2022). To assess the impact of the additional acquired RBD mutations on the stability of the domain, we used a differential scanning fluorimetry (DSF) assay that measures changes in protein intrinsic fluorescence as a thermal ramp is applied to the sample (**Figure 1C** and **S3**). The BA.1 RBD showed an inflection temperature (Ti) of 47.6 ± 0.2 °C, ∼7 °C lower than that of the WT RBD, similar to previous reports (Stalls et al., 2022). This loss in thermostability was partially recovered in the BA.2 RBD (Ti of 50.6 ± 0.1 °C). XBB.1.5 showed higher thermostability than the WT RBD with a Ti of ∼55.8 °C. Thus, mutations acquired in XBB.1.5 not only recovered the reduced thermostability observed in the early Omicron subvariants, but the XBB.1.5 RBD also surpassed the thermostability of the WT RBD. The RBDs of XBB.1.5 and XBB.1.16 differ by a single residue substitution at position 478. A T478K substitution that occurred in Omicron BA.1 was retained in BA.2, BA.5 and XBB.1.5, but was substituted to an Arginine in XBB.1.16 (**Figure 1A** and **S2**). As the DSF profiles of XBB.1.5 and XBB.1.16 RBDs were nearly identical (**Figure 1C**), we concluded that the K478R substitution, which is the only differing residue between the XBB.1.5 and XBB.1.16 RBDs, did not impact RBD thermostability. The EG.5 RBD showed a Ti of 52.5 ± 0.1 °C, which was ∼1.8 °C lower than that of the WT RBD and ∼3 °C lower than XBB.1.5 and XBB.1.16 RBD, implicating F456L, the only residue substitution between XBB.1.5 and EG.5, in the reduced stability of the EG.5 RBD.

Melting temperatures (Tm) of WT, XBB.1.5, XBB.1.6 and EG.5 RBD measured using Differential Scanning Calorimetry (DSC), showed the same thermostability trends as observed in the DSF experiments, i.e., EG.5 < WT < XBB.1.5 ∼XBB.1.16 (**Figure 1C** and **S3A**). The highest protein yields were obtained from the XBB.1.5 and XBB.1.16 RBD constructs, with both also being the most thermostable of the panel tested, while the BA.1 RBD was both the least thermostable and also gave the lowest in expression yield (**Figure S3B**).

In summary, RBD thermostability varied during SARS-CoV-2 Omicron evolution. A drop in thermostability observed with the Omicron BA.1 RBD, was progressively recovered in subsequent variants. XBB.1.5 and XBB.1.16 RBDs were more thermostable than the WT RBD, while EG.5 showed a reduction in thermostabilty compared to WT RBD, although it still remained more thermostable than BA.1, BA.2 and BA.5 RBDs. Stabilization of the RBD may lead to its more productive encounter with host receptor thereby improving virus transmissibility. Since the RBD is one of the most immunodominant regions of the S protein, its stability may also have vaccine implications, with improved stability resulting in a more durable immunogen, both in the S protein and RBD-only vaccine formats.

### ACE2 receptor binding of XBB lineage S protein ectodomains

To assess the impact of the acquired mutations on receptor engagement, we measured binding of the S protein ectodomains to the ACE2 receptor ectodomain by ELISA and Surface Plasmon Resonance (SPR) (**Figures 1D, 2A-G** and **S3C**). As expected based on previous studies (Yue et al., 2023, Uriu et al., 2023, Yamasoba et al., 2023), despite extensive mutation of the ACE2 binding region of the RBD, all the S proteins retained robust binding to ACE2. Indeed, the binding levels for all the Omicron S proteins tested were higher than for the D614G S protein, with XBB.1.5 showing the strongest ACE2 binding (**Figures 1D** and **2A**).

**Figure 2.**
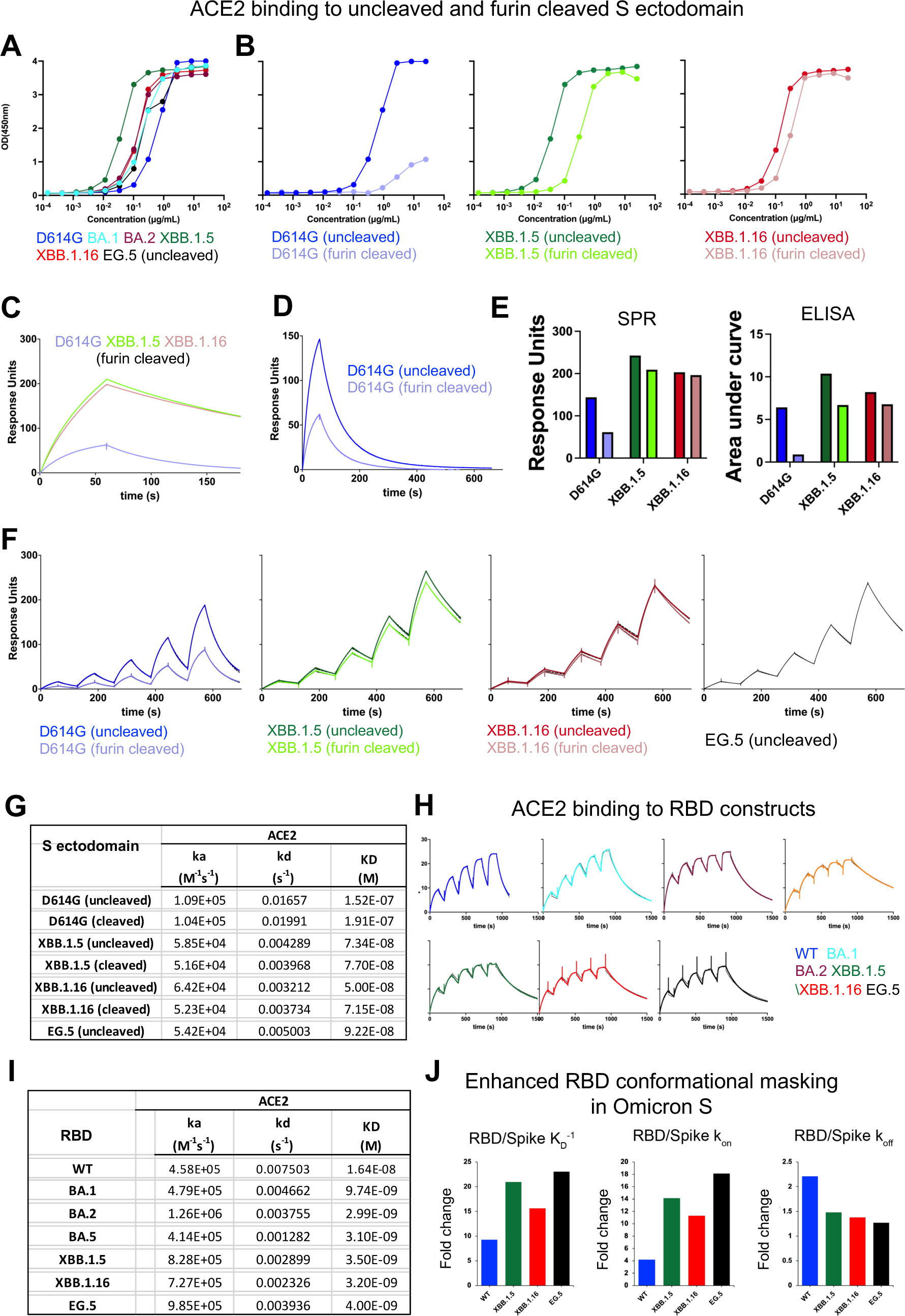
ACE2 binding to S ectodomains and RBD constructs, **A.** Binding of ACE2 to uncleaved S ectodomains measured by ELISA. **B.** Comparison of ACE2 binding to uncleaved and furin-cleaved S protein ectodomains of, from left to right, D614G (uncleaved shown in dark blue, furin-cleaved in light blue), XBB.1.5 (uncleaved in dark green, furin-cleaved in light green), XBB.1.16 (uncleaved in red and furin-cleaved in salmon). **C.** ACE2 binding to furin-cleaved S ectodomains analyzed by SPR using the same conditions as in Figure 1D. **D.** ACE2 binding to uncleaved and furin-cleaved D614G S ectodomains analyzed by SPR similarly as in 1D except with a longer dissociation time of 600s. **E.** Bar graph comparing the binding levels of ACE2 to uncleaved versus furin-cleaved S ectodomains. **F.** SPR single-cycle kinetics to measure ACE2 binding to S ectodomain constructs. **G.** Kinetics and affinity of ACE2 binding to the S ectodomains. **H.** ACE2 binding to RBD constructs measured by SPR using single cycle kinetics. **I.** Kinetics and affinity of ACE2 binding to the RBD constructs. The single cycle kinetics data were fit to a Langmuir 1:1 binding model. The data are shown in colored lines and the fit to the data are shown as overlaid black lines. **J.** Fold change in RBD versus S protein kinetics and affinities for ACE2.

With the S protein tethered to the sensor surface via its Twin-Strep tag, using SPR, we measured ACE2 binding affinities (or dissociation constant; K_D_) of 152 nM and 192 nM for the uncleaved (S-GSAS) and furin-cleaved (S-RRAR) D614G spikes, respectively (**Figures 2F-G**). ACE2 affinities for the uncleaved S ectodomains of XBB.1.5, XBB.1.16 and EG.5 were 73 nM, 50 nM, and 92 nM, respectively, while ACE2 affinity of furin-cleaved S ectodomain of XBB.1.5 and XBB.1.16 were 74 nM and 71 nM, respectively. EG.5 and EG.5.1 S ectodomains bound ACE2 with similar affinity and kinetics (**Figure S3E**). The tighter ACE2 affinity of the XBB.1.5, XBB.1.16, EG.5 and EG.5.1 S ectodomains compared to D614G was a result of the slower dissociation rates (k_off_) that also compensated for a reduction in association rates (k_on_) (**Figures 1D, 2C** and **2F-G**). ACE2 binding affinity and kinetics were similar for the uncleaved and furin-cleaved versions of the D614G, XBB.1.5 and XBB.1.16 S ectodomains. For the D614G variant, despite similar ACE2 binding affinities and kinetics, substantial and reproducible ACE2 binding level differences were observed between the furin-cleaved and uncleaved S proteins. The uncleaved D614G S protein showed higher levels of ACE2 binding compared to the furin-cleaved D614G S protein, suggesting reduced availability of ACE2-accessible “up” RBD in the furin-cleaved S protein compared to the uncleaved version. This observation is consistent with our previous reports of reduced RBD-up populations in furin-cleaved D614G S compared to the uncleaved S ectodomain (Gobeil et al., 2021a). By contrast, much less difference in binding levels was measured by SPR for the furin-cleaved versus uncleaved versions of the XBB.1.5 and XBB.1.16 S proteins, suggesting that the RBD up/down disposition in the XBB.1.5 and XBB.1.16 S proteins are less sensitive to furin-cleavage compared to the D614G S protein. These differences between the uncleaved and furin-cleaved S ectodomains were reflected also in the ACE2 binding levels measured by ELISA (**Figures 2A, B** and **E**).

In summary, and consistent with published reports (Yue et al., 2023, Uriu et al., 2023, Yamasoba et al., 2023), our data demonstrated higher affinity binding of the ACE2 ectodomain to SARS-CoV-2 S ectodomains of XBB.1.5, XBB.1.16, EG.5 and EG.5.1 variants than D614G, contributed by reduced dissociation rates. Further, differences in ACE2 binding levels observed for furin-cleaved versus uncleaved S ectodomains suggested a reduced impact of furin cleavage on domain dispositions of the XBB.1.5 and XBB.1.16 spikes than the D614G spike.

### ACE2 receptor binding of XBB lineage RBDs

Within the context of the S protein, the RBD is subject to conformational masking and can adopt an “up” or receptor accessible state where the RBM is exposed and available for binding to the host receptor, or a receptor inaccessible “down” state where the RBM is conformationally masked. To assess intrinsic binding of ACE2 to RBD without these conformational contraints, we measured ACE2 binding to a panel of RBD constructs derived from the D614G (or WT), BA.1, BA.2, BA.5, XBB.1.5, XBB.1.16 and EG.5/EG.5.1 S proteins (**Figures 2H-I** and **S3D**). The WT and XBB.1.5 RBD bound ACE2 with affinities of ∼16.4 nM and 3.5 nM, respectively, which are in close agreement with published studies (Irene et al., 2020, Lan et al., 2020, Walls et al., 2020, Yue et al., 2023). The ACE2 affinity for the Omicron RBDs were tighter than its affinity for the WT RBD, with BA.1 RBD binding ∼1.7-fold tighter and the other Omicron RBD constructs tested binding ∼4-5.5-fold tighter relative to WT RBD.

For variants where both the RBD and S ectodomians were tested, enhanced ACE2 affinities was observed for the RBD constructs than for the corresponding S proteins, with the fold change in RBD affinity with respect to S ectodomains affinity being higher for the Omicron XBB.1.5, XBB.1.16 and EG.5 variants compared to the D614G variant (**Figure 2I-J**). Higher association rates (k_on_) were primarily responsible for the enhanced ACE2 affinities in the RBD only constructs, consistent with a reduction in productive ACE2-RBM encounters when the RBD is part of the S protein. The differences in ACE2 binding affinity and kinetics between the RBD-only construct and the S ectodomains are a measure of the conformational masking that the RBM is subjected to in the context of the S protein. Our data demonstrate that conformational masking and shielding of the RBM is greater for the Omicron RBDs than the WT RBD. Thus, the increased ACE2 binding affinity of the Omicron S proteins relative to the D614G S protein (**Figure 2G**) was achieved by the increased intrinsic affinity of ACE2 for the Omicron RBDs counteracting the enhanced conformational masking of the RBM in the Omicron S proteins (**Figure 2H-J**).

### Cryo-EM structures of XBB lineage S ectodomains

To visualize their structures and conformations, we determined cryo-EM structures of S protein ectodomains of XBB.1.5, XBB.1.16, EG.5 and EG.5.1 (**Figures 3-5, Data S3-S4** and **Table S1**), utilizing our S-GSAS platform (**Figure S2** and **Data S1**) (Gobeil et al., 2021a). For XBB.1.16, we also determined the structure of S-RRAR-XBB.1.16, a construct that contained the native furin cleavage site between the S1 and S2 subunits, with the S protein cleaved with furin during protein expression (Gobeil et al., 2022) (**Figures 4**, **S2, Data S1, S3, S4** and **Table S1**).

**Figure 3.**
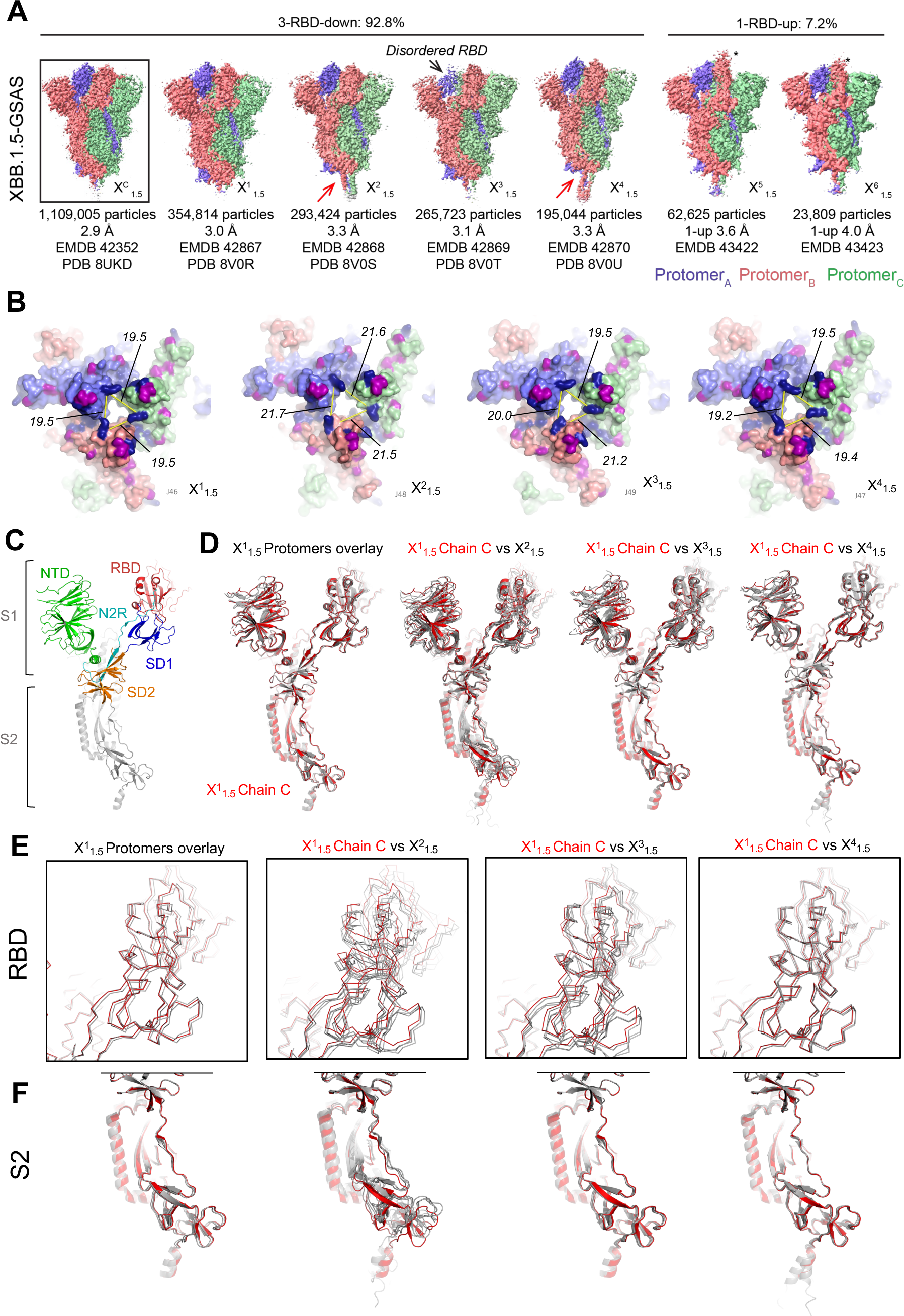
Cryo-EM structures of XBB.1.5 S protein ectodomains. **A.** Side views of cryo-EM reconstructions of the S-GSAS-XBB.1.5 ectodomain colored by chain. In the cryo-EM reconstructions of the 1-RBD-up states, the up positioned RBD in the map is identified by an asterisk. For the X^2^_1.5_ and the X^4^_1.5_ structures the position of the hinge at the C terminus is indicated by a red arrow. **B.** Tops views of the 3-RBD-down structures showing surface representation of the fitted coordinates, colored by chain. The RBD mutations that are shared between BA.2 and XBB.1.5 are colored in dark blue and the unique XBB.1.5 mutations are colored magenta. The distance between residue K440 in the three protomers for each structure are indicated (in Å). **C.** A single protomer of X^1^_1.5_ (chain C) with the S2 subunit colored grey and the S1 subunit colored by domain: NTD: green, N2R: cyan, RBD: red, SD1: blue, SD2: orange. **D.** Left to right. Overlay of the three protomers of X^1^_1.5_ with chain C colored red and the two other protomers colored grey. Overlay of the three protomers of X^2^_1.5_ with chain C of X^1^_1.5_ (colored red) and the three X^2^_1.5_ protomers colored grey. Overlay of the three protomers of X^3^_1.5_ with chain C of X^1^_1.5_ (colored red) and the three X^3^_1.5_ protomers colored grey. Overlay of the three protomers of X^4^_1.5_ with chain C of X^1^_1.5_ (colored red) and the three X^4^_1.5_ protomers colored grey. The SD2 subdomain (residues 592-697) was used for the superposition. **E.** Zoomed in view of the RBD from the superpositions shown in panel **D**. **F.** Zoomed in view of the S2 subunit from the superpositions shown in panel D.

**Figure 4.**
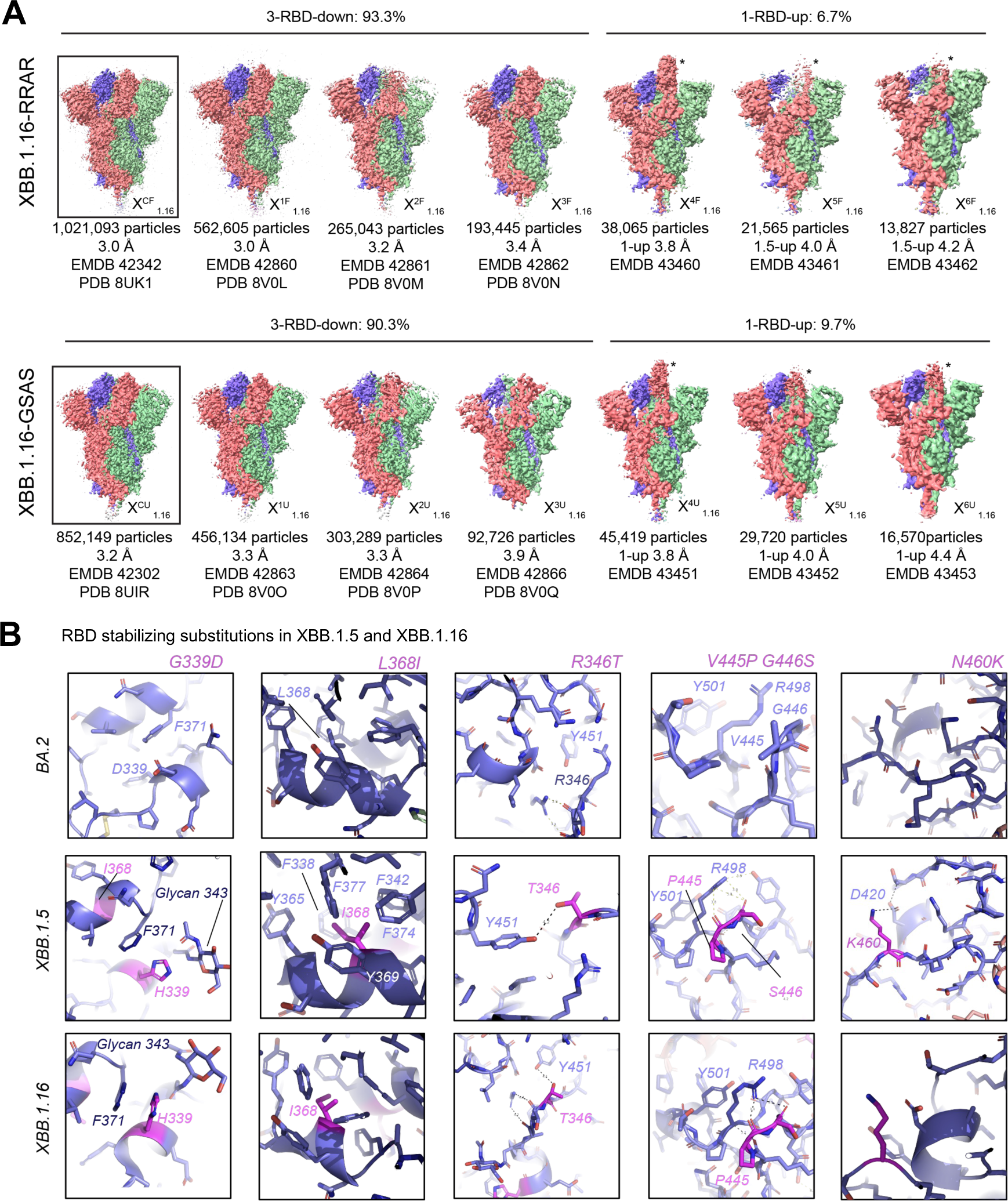
Cryo-EM structures of XBB.1.16 S protein ectodomains. **A.** Side views of cryo-EM reconstructions of the (top panel) S-GSAS-XBB.1.16 and (bottom panel) furin-cleaved S-RRAR-XBB.1.16 S ectodomains, colored by chain. In the cryo-EM reconstructions of the 1-RBD-up states, the up positioned RBD in the map is identified by an asterisk. **B.** Zoomed-in views of regions in BA.2 (top panel), XBB.1.5 (middle panel) and XBB.1.16 (bottom panel).

**Figure 5.**
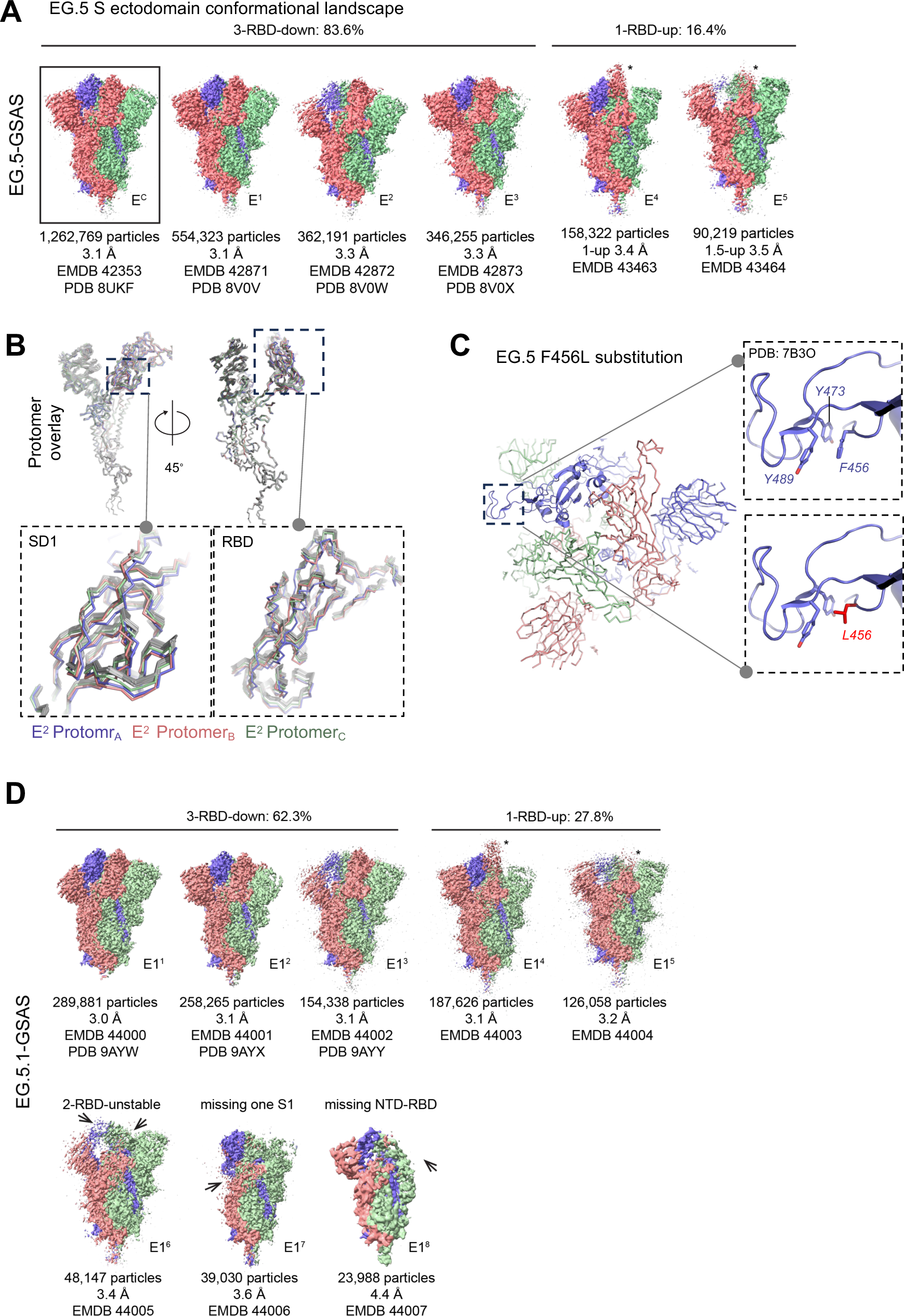
Cryo-EM structures of EG.5 and EG.5.1 S protein ectodomain, **A.** Side views of cryo-EM reconstructions of the S-GSAS-EG.5 S ectodomain colored by chain. In the cryo-EM reconstructions of the 1-RBD-up states, the up positioned RBD in the map is identified by an asterisk. **B.** Overlay of protomers of all EG.5 3-RBD down structures using S2 region 908-1035 for superposition. All protomers are colored grey except the protomers of the E^2^ structure that are colors salmon, blue and green. Insets show zoomed in views of the SD1 and RBD regions. **C.** The F456L substitution. The E1 structure is shown with a dotted box indicating the area that is zoomed in to show details of the environment around the F456L substitution. The detailed views are from PDB: 7B3O. **D.** Side views of cryo-EM reconstructions of the S-GSAS-EG.5.1 S ectodomain colored by chain.

#### Cryo-EM structures of the XBB.1.5 S protein ectodomain

For S-GSAS-XBB.1.5 (**Figure 3, Data S3-S4** and **Table S1**), we identified 3-RBD-down (closed) and 1-RBD-up (open) S ectodomain particle populations in a roughly 13:1 ratio of closed to open, which was substantially higher than the 3:1 and 2:1 ratios of closed to open S we had observed for BA.2 S and BA.1 S, respectively (Gobeil et al., 2022, Stalls et al., 2022). The dominant 3-RBD-down population yielded a 2.9 Å reconstruction, named X^DC^_1.5_ here (PDB: 8UKD; EMDB: 42352) (**Figure 3A**). Upon further subclassification, we obtained 4 and 2 classes, respectively, for the 3-RBD-down and 1-RBD-up populations. These were designated X^1^_1.5_ (PDB: 8V0R, EMDB: 42867), X^2^_1.5_ (PDB: 8V0S, EMDB: 42868), X^3^_1.5_ (PDB: 8V0T, EMDB: 42869), X^4^_1.5_ (PDB: 8V0U, EMDB: 42870), X^5^_1.5_ (EMDB: 43422), X^6^_1.5_ (EMDB: 43423).

In a study reporting the cryo-EM structural determination of XBB.1 S protein, only the 3-RBD-down, closed state of the S protein was identified in the cryo-EM dataset (Tamura et al., 2023), suggesting a strong propensity for adopting the 3-RBD-down closed state. Indeed, in this study initially the cryo-EM data analysis only yielded the 3-RBD-down state for the XBB.1.5 S ectodomain cryo-EM dataset, with the 1-RBD-up state being revealed upon extensive classification. The consensus 1-RBD-up population X^UC^_1.5_ showed a highly disordered up RBD characterized by missing or weak density (**Figure S4**). Upon further classification to resolve disorder of the X^UC^_1.5_ population, two subclasses, named X^5^_1.5_ and X^6^_1.5_ here, were revealed that differed primarily by the position of the up-RBD, with the RBD in X^6^_1.5_ adopting an intermediate conformation leaning more towards the central trimer axis and retaining contact with the RBD and NTD from the two adjacent protomers, while in X^5^_1.5_ the up RBD adopted a more erect conformation leaning away from the central trimer axis in one direction, and leaning sideways in an orthogonal direction towards the adjacent RBD and away from the adjacent NTD (**Figures 3A** and **S4**). Thus, the disordered or missing “up” RBD density in the consensus structure was a result of flexibility, with the RBD adopting varying dispositions around the canonical “up” configuration within the S protein molecules in the population.

Clear differences were discreible between the four XBB.1.5 3-RBD-down S protein subclasses (**Figure 3A-B**). The X^1^_1.5_ subclass consisted of the largest number of particles and most closely resembled the consensus structure X^DC^_1.5_ with an overall 0.26 Å C alpha root mean square deviation (Cα RMSD) between the two structures. The most divergent 3-RBD-down subclass was X^2^_1.5_ with overall 2.16 Å and 2.26 Å Cα RMSD, relative to X^DC^_1.5_ and X^1^_1.5_, respectively. The two other 3-RBD-down subpopulations X^3^_1.5_ and X^4^_1.5_ exhibited overall Cα RMSD of 0.32 Å and 0.37 Å, respectively, relative to the consensus X^DC^_1.5_, and 0.19 Å and 0.22 Å, respectively, relative to the X^1^_1.5_ subclass. Although not reflected in their overall Cα RMSDs, local differences were observed which distinguished the X^3^_1.5_ and X^4^_1.5_ populations from each other and from the other subclasses. In X^3^_1.5_, the cryo-EM map density for one of the three RBDs appeared weaker than the other two, suggesting variability in the RBD position. The residues at the C-termini in X^2^_1.5_ and X^4^_1.5_ were more ordered than they typically are in S ectodomain structures and were hinged at an angle relative to the body of the spike in a manner reminiscent of the hinging motion described in cryo-ET structures of the SARS-CoV-2 spike (Turonova et al., 2020). A notable difference observed between the 3-RBD-down subclasses were in the relative disposition of their RBDs, with the RBDs of two of the subclasses X^1^_1.5_ and X^4^_1.5_ more compactly packed than of the two others (X^2^_1.5_ and X^3^_1.5_) (**Figure 3B**). The separation between the RBDs was quantified by measuring the distance between the Cα atoms of residue K440 in the different protomers of each structure. All three distances were 19.5 Å for X^1^_1.5,_ and between 19.2 to19.5 Å for X^4^_1.5._ These distances measured between 21.5 to 21.7 Å for X^2^_1.5_ and between 19.5 to 21.2 Å for X^3^_1.5_.

The three protomers of X^1^_1.5_ overlaid by their SD2 subdomains showed good overlap throughout except for expected minor variations in the NTD region (**Figure 3C** and **D**), indicating a near symmetric structure despite no symmetry having been applied during the reconstructions. We next overlaid one of X^1^_1.5_ protomers with the three protomers of X^2^_1.5,_ X^3^_1.5_ and X^4^_1.5_, respectively. The RBD of the three X^4^_1.5_ protomers overlaid well with the RBD of X^1^_1.5_ (**Figure 3D** and **E**). The highest divergence of RBD positions between protomers was seen for X^2^_1.5_, followed by X^3^_1.5_. For X^3^_1.5_, although the RBD positions diverged, the S2 regions of the different protomers overlaid well with each other and with the S2 of X^1^_1.5_, while for the X^2^_1.5_ structure, differences in the S2 regions were visible throughout, both between its own protomers and with the X^1^_1.5_ protomer. In summary, the XBB.1.5 S ectodomain cryo-EM dataset revealed considerable diversity in its dominant 3-RBD-down population with conformational variability observed in RBD dispositions as well as in other regions of the S protein, including in the typically invariant S2 subunit.

#### Cryo-EM structures of the XBB.1.16 S protein ectodomain

The XBB.1.16 S protein differs from the XBB.1.5 S protein by two residue substitutions, E180V in the NTD and K478R in the RBD (**Figures 1A** and **S2**). The baseline forms of XBB.1.5 and XBB.1.16 showed similar ability to evade humoral immunity and therapeutic antibodies as did XBB.1, although both showed increased infectivity in pseudovirus assays compared to XBB.1 (Uriu et al., 2023, Yamasoba et al., 2023), with XBB.1.16 demonstrating greater growth advantage in the human population compared to XBB.1 and XBB.1.5 (Yamasoba et al., 2023).

In the S-GSAS-XBB.1.16 cryo-EM dataset (**Figure 4**, **Data S3-S4**, **Table S1**), we identified 3-RBD-down and 1-RBD-up S populations in a ∼9:1 ratio. We also determined the structures of the furin-cleaved S-RRAR-XBB.1.16 construct, where we observed ∼14:1 ratio of closed to open S conformations. The 1-RBD-up population of the uncleaved S-GSAS-XBB.1.16 S ectodomain showed expected variability in the disposition of the up RBD that was sampled in three subclasses, named X^4^_1.16_, X^5^_1.16_ and X^6^_1.16_. In the furin-cleaved S-RRAR-XBB.1.16 S ectodomain cryo-EM dataset, we identified not only 1-RBD-up S ectodomains but also populations where a second RBD appeared partially in the up position suggesting that the furin-cleaved S protein may be more flexible in the RBD-up or open states than the uncleaved XBB.1.16 S protein.

The 3-RBD-down structures of the uncleaved and furin-cleaved versions of the XBB.1.16 S ectodomain were similar with overall Cα RMSD of 0.3 Å. The 3-RBD-down population were subclassified into three populations each for the uncleaved and furin cleaved datasets of the XBB.1.16 S ectodomain. The populations were more homogeneous and the differences between the subclasses subtle compared to what was observed for the XBB.1.5 S protein, with overall Cα RMSDs between all the XBB.1.16 stuctures being in the 0.20-0.40 Å range.

Several residue substitutions in the XBB.1.5 and XBB.1.16 S proteins occurred at the ACE2 binding site (**Figure 1A**), including a rare F486P substitution that involved a two-nucleotide mutation and was implicated in the phenotypic change in XBB.1.5 that resulted in its gaining dominance over XBB.1. The F486P substitution facilitated enhanced ACE2 binding, greater transmissibility and escape from neutralizing antibodies (Uriu et al., 2023, Yue et al., 2023). F490S, another substitution in the RBM (residues 483–506), observed previously in the Delta VOC and Lambda VOI, was implicated in resistance to neutralizing antibodies (Tamura et al., 2023).

For XBB.1.5 and XBB.1.16 S ectodomains, we observed the RBD-RBD down state packing that we have previously reported for the BA.2 S protein (**Figure S5A**). No additional mutations were acquired at the RBD-RBD down state interprotomer contact sites of the XBB.1.5 and XBB.1.16 S proteins, and none of the previously acquired mutations at these sites were reverted or changed (**Figure 1A-B**). We found that the new RBD mutations either improved immune evasion by targeting the most immunodominant sites on the RBD or they facilitated interactions within the RBD, thereby improving RBD stability (**Figures 1A-C**). The Omicron BA.1 and BA.2 clades both carried a G339D substitution, and an additional S371F emerged in BA.2 that was preserved in subsequent linages. The G339D substitution is located at a site of intraprotomer RBD stabilization with the BA.2 S371F substitution facilitating close packing of two intra-RBD helical stretches 339–342 and 367–371 (**Figure 4B**). In XBB.1.5 a histidine appeared at residue 339 and further stabilized this region by packing against the side chain of F371 via a π-π stacking interaction, thus highlighting this region as a hotspot for RBD stabilization. Additionally, H339 was positioned to stack against and hydrogen bond with residue N343 and the N343 glycan, enhancing intra-helical stability and stabilizing and orienting glycan 343. Furthermore, the L368I substitution that appears in XBB.1.5 occurs within the 367-371 helical stretch at the center of an aromatic/hydrophobic cluster involving residues F371, Y365, F377, F374, F342, F338 and V367. Thus, it appears that the regions around the S371F, G339H and L368I are being optimized as Omicron evolves to stabilize the RBD while it engages in interprotomer down state RBD-RBD contacts.

Other stabilizing mutations in XBB.1.5 and XBB.1.16 include a R346T substitution in a RBD surface loop, a substitution that appeared in multiple lineages, including some that expanded, e.g. BQ.1.1, BA.4.6, BA.5.2.6, XBF, and FU.1. Shortening the side chain from an Arginine to a Threonine at position 346 brings the hydroxyl group within hydrogen bonding distance to the Y451 side chain hydroxyl (**Figure 4B**). A substitution of two consecutive residues V455P and G446S occurred in XBB.1.5 and XBB.1.16, where G446S was acquired in BA.1, reverted in BA.2 to G and changed to S again in XBB.1.5. Together they stabilize a turn in a surface loop that packs against residue R498 that we had previously shown forms a π-cation pair with Y501, both residues acquired in Omicron BA.1 via the Q498R and N501Y substitutions and retained in later variants. The R498-Y501 pair stabilizes a loop that contributes to interprotomer stabilizing contacts between down state RBDs. The N460K substitution occurs in a surface loop and forms a salt bridge with D420. In summary, the XBB lineage incorporated RBD substitutions that improved RBD stability while also maintaining the dominance of the 3-RBD-down state, which likely contributes to its immune evasive properties by conformational masking of the immunodominant RBD regions.

#### Cryo-EM structures of EG.5 and EG.5.1 S ectodomains

To visualize the structures and conformations of the EG.5 S protein, we obtained a cryo-EM dataset of S-GSAS-EG.5 (**Figures 5, Data S3-S4** and **Table S1**). EG.5 has a single F456L substitution relative to XBB.1.5 located in the RBD (**Figure 1A** and **S2**). The 3-RBD-down population still dominated in the EG.5 S cryo-EM dataset but its proportion relative the RBD-up populations (∼5:1) was smaller compared to what was observed in the XBB.1.5 and XBB.1.16 S ectodomain cryo-EM datasets. The EG.5 S protein 3-RBD-down consensus structure E^C^ (PDB: 8UKF; EMDB: 42353), was subclassified into E^1^ (PDB: 8VOV; EMDB: 42871), E^2^ (PDB: 8VOW; EMDB: 42872) and E^3^ (PDB: 8VOX; EMDB: 42873). Considerable variability between the 3-RBD-down subclasses was observed in the S1 subunit, with E^1^ being the most ordered with well-packed RBDs and E^2^ the least (**Figure S5B**). The three RBDs in E^2^ were asymmetrically arranged and showed variable densities with one RBD and its contacting NTD showing the weakest density indicating high disorder. Visualizing the Gaussian filtered map for this reconstruction confirmed that the down state assignment for this RBD. Similarly, one RBD-NTD pair was more disordered than the other two in E^3^, but less so than in E^2^.

Comparing the fitted coordinates for the 3-RBD-down structures showed that the subclass with the largest number of particles E^1^ was the most similar to the consensus structure E^C^ with an overall Cα RMSD of 0.30 Å, followed by E^3^ and E^2^ with Cα RMSDs of 0.48 and 0.64 Å, respectively, relative to E^C^. For all the structures, the S2 regions were very similar and overlaid well except at the C-terminal end where some conformational variability was observed (**Figure 5B**). The S1 regions of all except the E^2^ protomers were similar barring the positional fluctuations in the NTD region that are typically seen in SARS-CoV-2 S protein structures. For the E^2^ structure, two of the protomers showed shifted positions of the SD1 subunit and the RBD (**Figure 5B**). These shifts included the RBD of these protomers tilting away from the central vertical trimer axis, while still in the down position, and this RBD tilting in turn causing changes in the contacting NTD.

Since the resolution of the region around the EG.5 F456L substitution in the EG.5 cryo-EM structures precluded atomic level visualization, we relied on a crystal structure of the RBD (PDB: 7B3O) (Bertoglio et al., 2021) to understand the structural basis for the effects caused by the F456L substitution (**Figure 5C**). Residue F456 occurs in a region adjacent to the RBD hook that was named previously for its hook-like appearance, where F456 is part of an aromatic cluster with residues Y473 and Y489 (Gobeil et al., 2021b). The F456L substitution destabilizes this region by replacing the Phenylalanine with a less bulky, non-aromatic, albeit still hydrophobic residue. Since this region is a key epitope for neutralizing antibodies, the F456L induced structural changes likely contribute to its reported immune evasion properties (Jian et al., 2023). The destabilization of the F456/Y473/Y489 aromatic cluster due to the F456L substitution provides a structural rationale for the reduced thermostability of the EG.5 RBD relative to XBB.1.5 since this mutation is the only differing residue between the S proteins of these variants. F486L was recurrently selected for within the lineages we studied here subsequent to their initial expansion, indicating that despite its propensity to destabilize the RBD it conferred a selective advantage. It was preserved among EG.1 lineages, which originally emerged from within the XBB.1.9 lineage, but was also found independently in the XBB.1.16.6 sublineage, which was beginning to replace XBB.1.16 in North America prior to EG.1’s introduction, and the XBB.1.5 sublineages GK.1 (Pango alias XBB.1.5.70.1) and JD.1 (Pango alias XBB.1.5.102.1) which became dominant lineages in South America just as as EG.1 was expanding in other regions of the world (**Fig. S1**).

EG.5 acquired a single residue substitution Q52H in its NTD to become EG.5.1, the sublineage that spread most rapidly (Wang et al., 2023). Like EG.5, cryo-EM structures of the EG.5.1 S ectodomain (**Figure 5D and S5C**) showed heightened S1 subunit disorder, with an increased proportion of RBD up populations, as well small particle populations where parts of the NTD and RBD regions were disordered to an extent that these regions were not visible in the electron density. The disordered populations were reminiscent of similarly disordered structures observed in spikes that had transmitted between humans and minks (Gobeil et al., 2021b).

Taken together, our results reveal how substitution of one or two S1 subunit residues can have substantial local and global impacts on S protein structure. Placed in the context of the ongoing evolution of Omicron, our results show that different S protein properties are likely being optimized simultaneously. Therefore, despite a trend of increasing RBD stability during Omicron evolution, a single residue substition in EG.5 that destabilized the RBD was nevertheless selected for and retained in the subsequent expansion of the sublineage, likely due to its contribution to immune evasion at an immunodominant site.

### Antigenicity of the XBB lineage S proteins

To assess the impact of the acquired mutations on antigencity, we tested binding in ELISA to a panel of human anti-S antibodies that bind different regions of the S protein. These included S1 antibodies DH1050.1 (NTD-directed), DH1042 (RBD ACE2 binding site-directed), CR3022, DH1047 and S2X259 (RBD inner face directed), and DH1044, DH1193 and S309 (RBD outer face-directed) (**Figure 6A-C** and **S6**). We also included S2 antibodies DH1058.1 and DH1294 (fusion peptide-directed), DH1057.1 (stem helix epitope-directed), and a panel of Fab-dimerized glycan reactive antibodies that target a quaternary cluster of three S2 glycan moieties (**Figure 6D, F** and **S8**) (Li et al., 2021, Williams et al., 2021, Pinto et al., 2020, Yuan et al., 2020).

**Figure 6.**
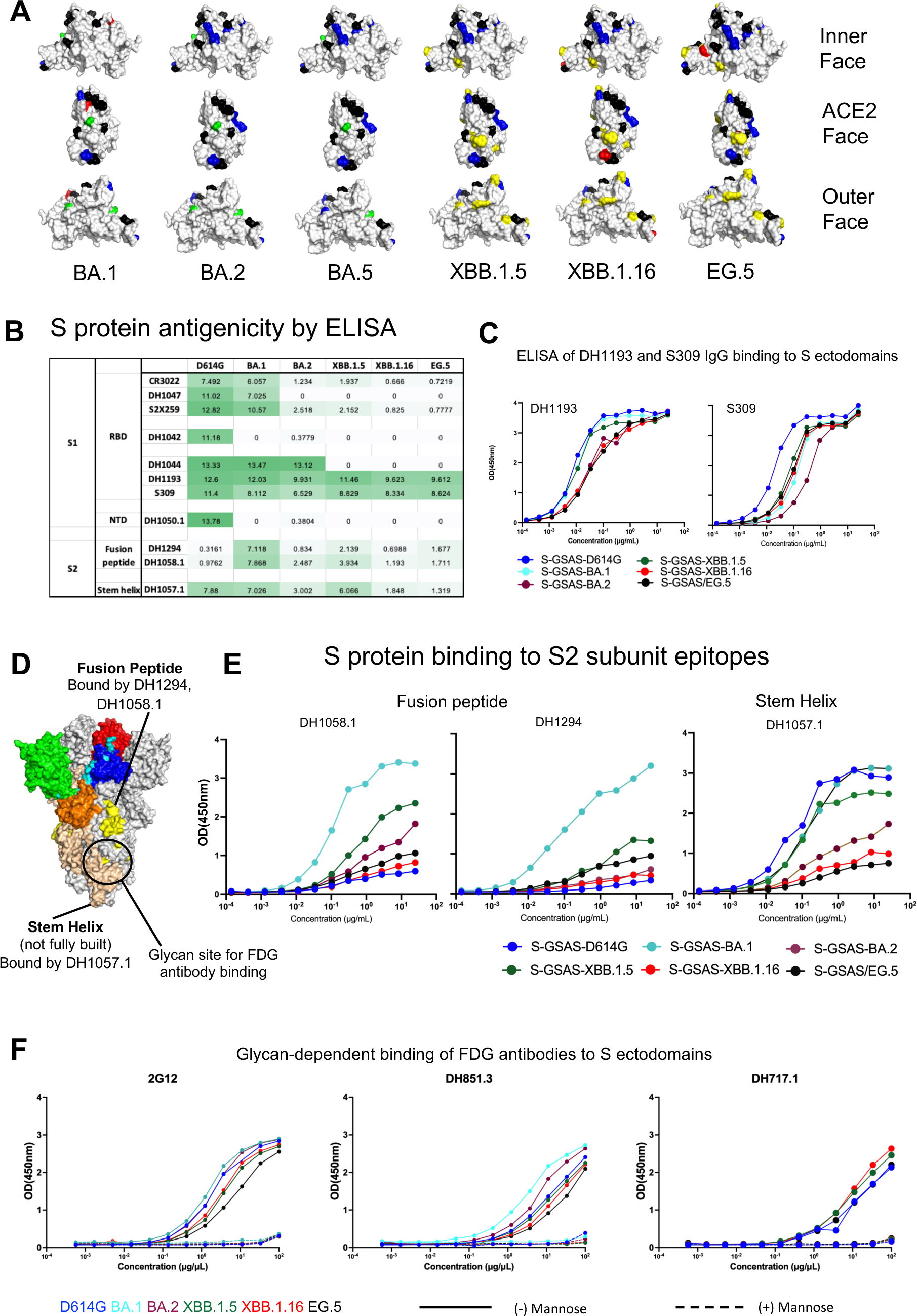
Antigenicity of XBB.1.5, XBB.1.16, and EG.5. **A.** Omicron mutations mapped to the RBD surface colored in grey. Color scheme followed is the same as in Figure 1A. Mutations that are common amongst all six variants are colored in black. Mutations that are shared by two or more are colored in blue. Mutations unique to only the BA.1, BA.2, BA.5 RBDs are colored in green. XBB.1.5, XBB.1.16, and EG.5 unique mutations are in yellow. A mutation unique to only one of the RBDs is colored in red. **B.** ELISA Binding of D614G, BA.1, BA.2, XBB.1.5, XBB.1.16, and EG.5 S ectodomains to antibodies targeting different regions of the S protein. Binding was quantified using LogAUC values of the ELISA curves, and color coded on a scale of dark green to white with darker green indicating stronger binding, lighter green indicating weaker binding and white indicating no binding. **C.** ELISA binding of RBD outer face directed antibody DH1193 to S ectodomains. **D.** Surface representation of a SARS-CoV-2 S protein structure with two protomers colored gray and one colored by domains; NTD: green, N2R: cyan; RBD: red, SD1:blue. SD2: orange. The fusion peptide (DH1058.1 and DH1294), stem helix (DH1057.1) and glycan cluster (FDG antibodies) epitopes are labeled. **E.** Binding of S ectodomains to S2 subunit binding antibodies; DH1058.1 and DH1294 (fusion peptide directed) and DH1057.1 (stem helix epitope directed). **F.** ELISA Binding of FDG antibodies 2G12, DH851.3 and DH717.1 to S ectodomains, either in absence (solid line) or presence (dashed line) of 1M Mannose.

Of all the S1 directed antibodies tested, the only two that showed appreciable binding to the XBB.1.5, XBB.1.16 and EG.5 S ectodomains in ELISA were DH1193 and S309 that bind distinct, albeit overlapping, epitopes on the RBD outer face (**Figure 6A-C, S6**) (Gobeil et al., 2022). The ELISA binding of DH1193 and S309 translate to their ability to neutralize in pseudovirus neutralization assays (Case et al., 2022, Wang et al., 2023, Iketani et al., 2022, Malewana et al., 2023, Stalls et al., 2022). In a recent study, neutralizing antibodies that bound the DH1193 epitope region have been elicited in non-human primates by vaccination, highlighting this region of the RBD as a vaccine target for eliciting neutralizing antibodies that may be less resistant to mutations acquired during SARS-CoV-2 evolution (Malewana et al., 2023). The EG.5 and EG.5.1 S ectodomains showed similar antigenic properties with robust binding retained to the RBD outer face targeting antibodies S309 and DH1193 (**Figure S7**).

At the S2 subunit, a panel of Fab-dimerized glycan reactive (FDG) antibodies that target a quaternary S2 glycan cluster bound XBB.1.5, XBB.1.16 and EG.5 S ectodomains in a glycan dependent manner (**Figure 6F**). Consistent with previous observations, we found that the furin-cleaved S ecotodomains showed stronger binding to the FDG antibodies than the uncleaved versions, demonstrating allosteric effect of furin cleavage on S2 structure and on the presentation of the glycan epitope (**Figure S8**).

Structural modeling showed that the epitopes of the fusion peptide targeting antibodies DH1058.1 and DH1294, and stem helix targeting antibody DH1057.1, are not fully accessible for antibody binding on the pre-fusion SARS-CoV-2 S protein (Gobeil et al., 2022, Kapingidza et al., 2023). Despite this, measurable binding of the SARS-CoV-2 S protein to these antibodies can be observed by SPR and ELISA, suggesting antibody-induced local refolding of the S protein structure to expose the antibody eptiopes. For the fusion peptide-directed antibodies, we have previously reported increased ELISA binding for the BA.1 S ectodomain relative to the D614G S ectodomain, and interpreted this to indicate increased flexibility of this region in the BA.1 S protein leading to greater access to the fusion peptide in the presence of the antibody (Stalls et al., 2022, Gobeil et al., 2022). BA.2 S showed reduced binding to DH1058.1 and DH1294 in ELISA, relative to BA.1 S, indicating stabilization of this region in the pre-fusion S conformation (Stalls et al., 2022). Our results here recapitulate these trends with BA.1 S showing the highest levels of binding, D614G S the lowest and BA.2 S intermediate between the two (**Figure 6E**). XBB.1.16 and EG.5 S proteins show lower binding to DH1058.1 than BA.2 S, while XBB.1.5 shows higher binding than BA.2 S, although still substantially lower than BA.1 S, demonstrating overall stabilization of this S protein region in its pre-fusion conformation upon Omicron evolution, after being destabilized in the first identified Omicron variant BA.1. The difference in ELISA binding observed between XBB.1.5 and XBB.1.16 is notable and consistent with the greater diversity of S2 subunit conformation observed in the XBB.1.5 cryo-EM subpopulations, suggesting greater S2 conformational malleability in the XBB.1.5 S protein than in XBB.1.16 S. The second fusion peptide-directed antibody tested DH1294 shows similar trends, although the differences were less pronounced than they were with DH1058.1 (**Figure 6E**).

The trends observed for DH1057.1 (stem helix) binding was distinct from those observed with the fusion peptide-directed antibodies, with strongest binding observed for the D614G, BA.1 S and XBB.1.5 S ectodomains, and BA.2, XBB.1.16 and EG.5 S ectodomains binding at substantially lower levels (**Figure 6E**). The differences in the observed trends between the fusion peptide directed antibodies and the stem helix antibody suggest differences in the mechanisms of exposure of their epitopes. The substantially stronger binding observed for XBB.1.5 compared to XBB.1.16 or EG.5 further demonstrates the allosteric effect of one or two S1 subunit mutations on the S2 subunit stem-helix epitope.

Taken together, our antigenicity data reveal how differently S1 and S2 subunit epitopes have been impacted by SARS-CoV-2 evolution. Widespread loss in binding was observed to S1-directed antibodies, leaving only selected epitopes such as those on the RBD outer face that bind to antibodies like S309 and DH1193 still capable of binding antibodies elicited against the initial Wu.1 spike. S2 subunit epitopes have also been affected in large part due to conformational effects emanating from S1 subunit residue chnages. Despite a difference of only two S1 subunit residues between the XBB.1.5 and XBB.1.16 S proteins, the differences in their binding to the fusion peptide-directed and stem helix loop directed antibodies demonstrate allosteric effects of the S1 residue substitution on the S2 subunit. For the XBB.1.5 and XBB.1.16 S proteins, these differences in S2 subunit antibody binding were consistent with the differences in the S2 subunit observed in their cryo-EM populations (**Figure 3-4**). Similarly, despite a single RBD residue substitution between XBB.1.5 and EG.5 S proteins, the two ectodomains show distinct binding profiles to the fusion peptide-directed and stem helix-directed antibodies.

### Vector analysis

We next utilized a previously described interprotomer vector-based analysis to examine the Omicron sub-lineage S protein domain arrangements (Henderson et al., 2020) (**Figure 7**). This analysis generates vectors between domains across each chain using domain centroids and anchor residues to define the quaternary geometry of the S protein domains (**Figure 7A**). We showed previously that this geometry differed markedly between MERS, SARS-CoV-1 and SARS-CoV-2 and that these geometries have shifted as SARS-CoV-2 has evolved (Gobeil et al., 2022, Henderson et al., 2020, Stalls et al., 2022). We used these vectors to calculate domain-to-domain distances, angles and dihedrals to determine which vary the most using principal components analysis (PCA). The dataset composed of vectors from the Omicron sub-lineage S ectodomain structures determined here, and our previously determined structures for the D614G, mink, Alpha, Beta, Delta variants and the K417N-E484K-N501Y triple mutant (TM) that appeared in Beta and Gamma VOCs, showed a distinct “Omicron cluster” in the first principal component (**Figure 7B**). This cluster showed substantially less variability than the earlier variants in both the first and second principal components. Examination of the vector-based contributions to the first principal component indicated the primary driver of differences between earlier variants and the Omicron variant and sub-lineages were (1) the dihedral orientations between the RBDs, (2) the RBD-to-RBD distances, and (3) the Subdomain 1 (SD1) to S2-central helix distances.

**Figure 7.**
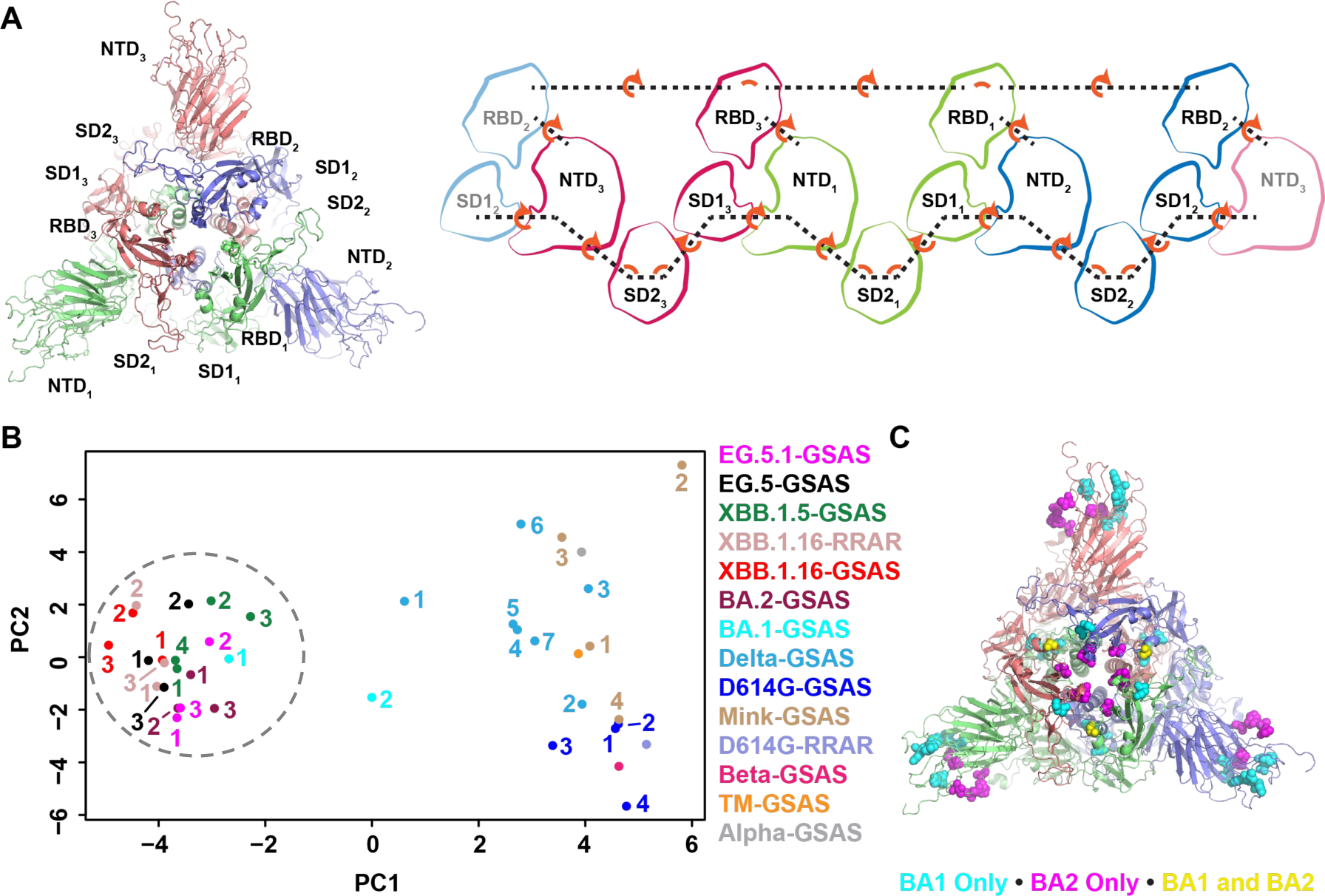
Mutations in the SARS-CoV-2 Spike RBD-to-RBD interfaces create a unique, stable down state domain arrangement in Omicron sub-lineages. **A.** (left) Top-down view of the SARS-CoV-2 Spike highlighting the positions the S1 subunit domains. (right) A “rolled-out” two-dimensional representation of the domains showing the interprotomer vectors (dashed lines) and the angles and dihedrals between them. **B.** SARS-CoV-2 variant Spike interprotomer vector principal components analysis. Each point represents a single down state Spike trimer. PDB IDs [D614G: 1, 7KE4; 2, 7KE6; 3, 7KE7;4,7KE8], [Alpha: 7LWS], [Beta: 7LYL], [TM (Triple Mutant spike with K417N, E484K and N501Y mutations): 8CSA], [Delta: 1, 7TOX; 2, 7TOY; 3, 7TOZ; 4, 7TPO; 5, 7TP1; 6, 7TP2; 7, 7TOU], [Mink: 1, 7LWI; 2, 7LWJ; 3, 7LWK; 4, 7LWL], [BA.1: 1, 7TL1; 2, 7TF8], [BA.2: 1, 7UB0; 2, 7UB5; 3, 7UB6], [XBB.1.5-GSAS: 1, 8V0R; 2, 8V0S; 3, 8V0T; 4, 8V0U], [XBB.1.16-RRAR: 1, 8V0L; 2, 8V0M; 3, 8V0N], [XBB.1.16-GSAS: 1, 8V0O; 2, 8V0P; 3, 8V0Q], [EG.5-GSAS: 1, 8V0V; 2, 8V0W; 3, 8V0X]. The dashed circle indicates the “Omicron cluster”. **C.** (left) Side view of the SARS-CoV-2 Spike with BA.1 and BA.2 mutation residues shown as spheres. Mutations colors are according to positions that are only mutated in BA.1 (cyan), BA.2 (magenta), or that are mutated in BA.1 and BA.2 but for which the mutations differ (yellow). (right) Top-down view of the Spike showing the positions of the mutations in the RBDs and NTDs.

A single BA.1 structure (formerly named O^1^_BA.1_; PDB: 7TF8) (Gobeil et al., 2022) that we previously determined appeared intermediate to the two primary principal component clusters separating the Omicron and earlier variants, while the BA.2 variant structures resided exclusively in the Omicron cluster. This difference suggests mutations that differ between BA.1 and BA.2 cause the Omicron sub-lineages to settle into a stable domain geometry that was retained in subsequent variants. To determine the driving factor in this reduction in domain-to-domain variability in the Omicron sub-lineages, we compared mutation positions between BA.1 and BA.2 that either occurred only in BA.1, only in BA.2, or occurred in both BA.1 and BA.2 but differed in identity (**Figure 7C**). While the BA.1 SD1 T547K mutation may have affected the structure, the BA.2-only RBD-to-RBD interface mutations T376A, D405N, and R408S stood out as likely drivers of the difference between BA.1 and BA.2 that led to the stabilized domain geometry. Taken together, the results from our vector analysis show that close coupling of the RBD domains in the Omicron sub-lineages creates a sterically restricted S1 domain.

## Discussion

The emergence of the Omicron variants marked a dramatic inflection point in the COVID-19 pandemic. The primary characteristic that distinguished Omicron from previous variants was its substantially improved immune evasion. While the pandemic has ended as public health risk has declined, Omicron remains a topic of public health concern as new immune evasive subvariants continue to emerge, necessitating continued surveillance and updated vaccine boosters. The most updated booster currently is a monovalent vaccine based on the XBB.1.5 S protein. In this study, we have characterized the structure, receptor binding and antigenicity of the XBB.1.5 S ectodomain, along with the S ectodomains of three XBB lineage members, XBB.1.16, EG.5 and EG.5.1, that followed it. Our data show that the XBB.1.5 RBD is more stable compared to the RBDs of previous variants, as well as the RBD of the EG.5/EG.5.1 sublineage that dominated a subsequent global sweep. A more durable RBD may bode well for the current booster. Additionally, compared to the other Omicron variants tested, the XBB.1.5 S protein showed increased antibody accessibility to subdominant S2 subunit epitopes, particularly at the stem helix region, which may better enable elicitation of potentially protective antibodies that target these regions (Pinto et al., 2021, Kapingidza et al., 2023).

Our structural studies of the early Omicron S protein variants BA.1 and BA.2 have revealed accumulation of mutations leading to stabilizing interprotomer interactions between down state RBDs, resulting in a tendency to favor the 3-RBD-down conformation that hold the RBD in a closed, receptor inaccessible conformation (Gobeil et al., 2022, Stalls et al., 2022). This trend, first observed in BA.1, was further fine-tuned in BA.2, where additional mutations optimized the RBD-RBD down state packing, while also stabilizing the RBD itself. This was in stark contrast to previous VOCs with a tendency to favor RBD-up conformations, indicating that among earlier variants transmissibility was a dominant selective force. This shift allowed SARS-CoV-2 to remain relevant in an environment where considerable population level immunity had been achieved through vaccination, natural infection or both. The XBB lineage S ectodomains studied here have retained the overall domain organization of the BA.2 S protein, with Omicron S protein structures forming a distinct cluster in our vector analysis that is well-separated from pre-Omicron VOCs (**Figure 7**). The vector analysis highlights the BA.1 S protein architecture to be a transition point with one BA.1 S ectodomain structure residing within the Omicron cluster while another appeared intermediate between the Omicron and pre-Omciron distributions. This pattern of progressive transition of the S protein architecture is also demonstrated in Difference Distance Matrix (DDM) analysis, which provides a superposition-free way to compare two structures (**Data S5-S6)**. The XBB.1.5, XBB.1.16 and EG.5 S ectodomains are, thus, more similar to the BA.2 S protein, less so to the BA.1 S and least similar to the D614 G S protein.

The RBD “down” conformation hides the most immunodominant and conserved RBD regions, making it an effective immune evasion strategy. The stabilization of this 3-RBD-down state, however, presents an apparent conundrum since the RBD-up conformation is rquired for receptor binding and transmission. How does Omicron maintain high affinity receptor binding and transmissibility while retaining and improving the occupancy of its 3-RBD-down state? For the early Omicron BA.1 variant, we had described a strained structure that we envisioned was primed to transition to an open conformation in the presence of receptor (Gobeil et al., 2022). Our structural conclusions were supported by subsequent HDX-MS studies that revealed a BA.1 spike more readily able to transition to an open state in the presence of the ACE2 receptor (Calvaresi et al., 2023). As Omicron evolved from BA.1 to BA.2, it appeared to have resolved some of the structural strain, although the conformational diversity observed with the XBB.1.5 S 3-RBD-down population, including conformational varibility in the S2 subunit, suggests that internal destabilization or pre-priming may still be a parameter that the evolving Omicron lineage is continuing to optimize. A second factor that enabled Omicron to maintain high affinity receptor binding was the evolution of the RBD itself with mutations within the domain stabilizing the RBD as well as enhancing its intrinsic affinity for ACE2 (Yue et al., 2023, Uriu et al., 2023). Finally, a more electropositive host membrane facing surface may have facilitated Omicron S protein interactions with host cell surface coreceptors such as heparan sulfate to increase accumulation of the virus near the host cell and facilitate spike opening (Clausen et al., 2020, Kearns et al., 2022, Kim et al., 2023). An increased electropositive surface on the host membrane facing side was noted for the Omicron BA.1 S protein (Gan et al., 2022). For the XBB lineage structures determined here, although there is an overall reduction in positive potential across the surface due to residue subsitutions, the positive electrostatic potential at the center of the host facing surface of the spike is maintained (**Figure S9**). Indeed, the Omicron variant may be taking a page out of the playbook of the currently endemic coronaviruses that typically adopt the 3-RBD-down conformation with interactions with the host glycocalyx facilitating the transition of the RBD to the up position, thus enabling optimal immune evasion while ensuring the required transition to the conformation that can bind host receptor (Pronker et al., 2023).

The ongoing evolution of the Omicron variant spikes involves multi-parameter optimization where the major driver is immune evasion. The other properties that need to be simultaneously optimized include pre-fusion conformation stability, host receptor binding and ability to undergo the conformational changes required for entry into host cell. Our study shows how the XBB lineage gained increased immune evasive properties while improving spike stability and receptor binding, and how it has continued to evolve. The substantial structural, conformational and antigenic impact that were effected by only one or two residue substitutions underscored the effectiveness of allosteric networks in the SARS-CoV-2 S protein and foretell its continued evolution to escape host immunity.

## Acknowledgements

Cryo-EM data were collected at the Duke Krios at the Duke University Shared Materials Instrumentation Facility (SMIF), a member of the North Carolina Research Triangle Nanotechnology Network (RTNN), which is supported by the National Science Foundation (award number ECCS-2025064) as part of the National Nanotechnology Coordinated Infrastructure (NNCI). This study utilized the computational resources offered by Duke Research Computing (http://rc.duke.edu; NIH 1S10OD018164-01) at Duke University. This work was supported by NIH R01 AI165947 (P.A. and R.H.), R01 AI 165147 (P.A. and W.B.W.), NIH, NIAID, DMID grant P01 AI158571 (B.F.H.).

## Author contributions

P.A.; Methodology – P.A. and R.H; Investigation –R.Parsons, B.T., R.Parks, M.M., C.S.P., X.H., S.S., K.J., S.M-C.G., R.J.E., D.C.M., B.K.; Data Curation – Q.E.Z., J.J.L., C.S.P., X.H., S.S., K.J., S.M-C.G., R.J.E., R.H. and P.A.; Validation – Q.E.Z., J.J.L., R.H., and P.A.; Resources – K.O.S. and P.A.; Visualization – Q.E.Z., J.J.L., R. Parsons., R.H., and P.A.; Writing - Original Draft – Q.E.Z., J.J.L., R.Parsons. R.H. and P.A.; Writing - Review & Editing – all authors; Supervision – R.J.E, W.B.W., B.F.H. and P.A.; Project Administration –P.A.; Funding acquisition – W.B.W., B.F.H., R.H., and P.A.

## RESOURCE AVAILABILITY

### Lead Contact

Further information and requests for resources and reagents should be directed to and will be fulfilled by the Lead Contact, Priyamvada Acharya.

### Materials Availability

Further information and requests for resources and reagents should be directed to Priyamvada Acharya. Plasmids generated in this study have been deposited to Addgene with the following accession numbers: 212990, 212991, 212992, 212993, 213037, 213038, 213070, 213410, 213478.

### Data and Code Availability

- Cryo-EM reconstructions and atomic models generated during this study are available at wwPDB and EMBD (https://www.rcsb.org; http://emsearch.rutgers.edu) under the accession codes PDB IDs 8UKD,8UIR, 8UK1, 8UKF, 8V0L, 8V0M, 8V0N, 8V0O, 8V0P, 8V0Q, 8V0R, 8V0S, 8V0T, 8V0U, 8V0V, 8V0W, 8V0X, 9AYW, 9AYX, 9AYY and EMDB IDs 42302, 42342, 42352, 42353, 42860, 42861, 42862, 42863, 42864, 42866, 42867, 42868, 42869, 42870, 42871, 42872, 42873, 43422, 43423, 43451, 43452, 43453, 43460, 43461, 43462, 43463, 43464,44000, 44001, 44002, 44003, 44004, 44005, 44006, 44007.
- This paper does not report original code.
- Any additional information required to reanalyze the data reported in this paper is available from the lead contact upon request.

## METHOD DETAILS

### Plasmids

The plasmids in this study were obtained by site-directed mutagenesis, performed by GeneImmune Biotechnology (Rockville, MD) with quality controls. The S protein sequences are adapted from residues 1 to 1208 (GenBank: MN908947) with a D614G mutation, furin cleavage site (RRAR; residue 682-685) retained or mutated to GSAS. All constructs are attached to a C-terminal T4 fibritin trimerization motif, a C-terminal HRV3C protease cleavage site, a TwinStrepTag and an 8XHisTag, sequentially, and incorporated into pαH vector. The RBD constructs were originated from BEI resources (Catalog number NR-52309). The coding region was codon optimized and subcloned into pCAGGS vector for mammalian expression. All plasmids in this study are deposited to Addgene (https://www.addgene.org).

### Protein Expression and Purification

Plasmids encoding the S protein ectodomains were transfected into Gibco FreeStyle 293-F cells (embryonal, human kidney) following the manufacturer’s recommendations. Supernatant was harvested 6 days post transfection and filtered through a 0.22 µm filter. StrepTactin resin (IBA LifeSciences) was used to purify the S ectodomains, followed by size exclusion chromatography (SEC) on s <u>Superose</u> 6 10/300 GL Increase column (Cytiva, MA) in 2 mM Tris, pH 8.0, 200 mM NaCl, 0.02% NaN_3_. Quality check of the proteins was completed by NuPage 4-12% (Invitrogen, CA) SDS-PAGE. SEC fractions containing the S protein were combined and concentrated. The final products were flash-frozen in liquid nitrogen and stored at −80°C in single-use aliquots for future use. Aliquots were thawed at 37 °C for 20 minutes before use.

Spike ectodomains were harvested from the concentrated supernatant on day 6 post transfection. The spike ectodomains were purified using StrepTactin resin (IBA LifeSciences) and size exclusion chromatography (SEC) using a Superose 6 10/300 GL Increase column (Cytiva, MA) equilibrated in 2 mM Tris, pH 8.0, 200 mM NaCl, 0.02% NaN3. All purification steps were performed at room temperature within a single day. Protein quality was assessed by SDS-PAGE using NuPage 4-12% (Invitrogen, CA). The purified proteins were flash frozen and stored at −80°C in single-use aliquots. Each aliquot was thawed by a 20-min incubation at 37°C before use.

Wild-type RBD and RBD variants were produced in 293F cells and harvested from supernatant on the 6th day post transfection. The RBDs containing an 6x-Histidine tag were purified via nickel affinity chromatography using a HisTrap excel column (Cytiva, MA). Supernatant was loaded onto the column and the column washed with buffer A (1x PBS pH 8.0) until baseline. The proteins were eluted from the column by applying a gradient over 20 CV from 100% buffer A to 100% buffer B (1x PBS pH 8.0, 1 M Imidazole). Fractions containing RBD were pooled, concentrated, and further purified by size exclusion chromatography (SEC) using a Superdex 200 Increase 10/300 GL column (Cytiva, MA) equilibrated with 1x PBS, pH 8.0. All steps of the purification were performed at room temperature. Protein quality was assessed by SDS-PAGE using NuPage 4-12% (Invitrogen, CA). The purified proteins were flash frozen and stored at −80°C in single-use aliquots. Each aliquot was thawed at 4°C before use. ACE2 containing a 6x-Histidine tag was prepared similarly to RBD except size exclusion chromatography was performed using a Superose 6 10/300 GL column.

Antibodies were produced in Expi293F cells and purified by Protein A affinity. Fab fragments were generated by digestion of the antibodies using LysC. Antibodies and Fabs were further purified using SEC using a Superose 6 Increase 10/300 GL column (Cytiva, MA).

### Differential scanning calorimetry

DSC assays were performed using an Affinity NanoDSC (TA Instruments, DE). RBDs were diluted to approximatively 3 mg/mL. Heat (uW) was measured during the protein melting over a temperature range from 10 – 100 °C. Data were processed and fitted using a two-state scaled model with the NanoAnalyze software version 3.12.5 (TA Instruments, DE).

### Differential scanning fluorimetry

DSF assays were performed using Tycho NT.6 (NanoTemper Technologies). RBDs were diluted to approximatively 1 mg/mL. Intrinsic fluorescence was measured at 330 nm and 350 nm while the sample was heated from 35 to 95°C at a rate of 30°C/min. The ratio of fluorescence (350/330 nm) and inflection temperatures (Ti) were calculated using the inbuilt software in the Tycho NT. 6.

### ELISA assays

384-well plates were coated with 2 µg/ml of streptavidin (Thermo Fisher Scientific, MA), followed by blocking. Spike ectodomains at 2 µg/ml were applied to the plates and incubated at room temperature for 1 hour, following which the spike solution was removed and the wells washed. The mAbs and ACE2 were serial diluted from 25 µg/ml with 3-fold each dilution. Various concentrations of mAbs or ACE2 were added to corresponding wells and incubated at room temperature for 1 hour, following which excess solution was removed and the wells were washed. Goat anti-human IgG HRP (1:15000) and the substrate 3,3′,5,5′-tetramethylbenzidine (TMB) (Sera Care Life Sciences no. 5120-0083) were used to detect binding with an incubation time of 15 minutes. Absorbance was measured at 450 nm. The binding levels were quantified by log area under the curve (Log_AUC_).

SARS-CoV-2 S ectodomains were tested for binding to recombinant FDG mAbs in ELISA in the absence or presence of single monomer D-mannose as previously described (Williams et al., 2021). Briefly, 20 ng spike proteins were captured by streptavidin (30ng per well) to individual wells of a 384-well Nunc-absorb ELISA plates using PBS-based buffers and assay conditions as previously described (Williams et al., 2021, Alam et al., 2017, Bonsignori et al., 2017). D-mannose (Sigma, St. Louis, MO) was used to compete mAb binding to glycans on the S proteins; D-mannose solutions were also prepared in ELISA PBS-based glycan buffers at a concentration of [1M] D-mannose as described (Williams et al., 2021). Mouse anti-monkey IgG-HRP (Southern Biotech, CAT# 4700-05) and Goat anti-human IgG-HRP (Jackson ImmunoResearch Laboratories, CAT# 109-035-098) secondary antibodies were used to detect antibody bound to the S proteins. HRP detection was subsequently quantified with 3,30,5,50-tetramethylbenzidine (TMB) by measuring binding levels at an absorbance of 450nm, and binding titers were also reported as Log area under the curve (AUC).

### Surface Plasmon Resonance

Binding experiments were performed using SPR on a Biacore T-200 (Cytiva, MA) with HBS buffer supplemented with 3 mM EDTA and 0.05% surfactant P-20 (HBS-EP+, Cytiva, MA). All binding assays were performed at 25 °C.

ACE2 binding to Spike proteins was assessed using a Series S SA chip (Cytiva, MA). Spike were coated on the surface at 200 nM (120s at 5 µL/min. For single injection experiments, ACE2 was injected at 200 nM over the chip surface (60s at 50 µL/min) with an off time of 120s. The surface was regenerated three pulses of a 1 M NaCl and 50 mM NaOH solution for 10 seconds at 100µL/min. Kinetics experiments were performed using single cycle kinetics mode with five different concentrations of ACE2 serially diluted two-fold. ACE2 concentrations ranged from 12.5 – 200 nM. Injections lasted 60 sec with a final off time of 120 seconds at 50 µL/min.

ACE2 binding to RBDs was performed via amine coupling of the ACE2 to the chip surface. ACE2 was coupled to a Series S CM5 sensor chip (Cytiva, MA) using standard amine coupling chemistry to approximately 800 RU. For single injection experiments, RBDs were injected at 100 nM (120s at 50 uL/min) with an off time of 20 minutes. Kinetics experiments were performed using single cycle kinetics mode with five different concentrations of each RBD serially diluted two-fold. RBD concentrations ranged from 12.5 – 200 nM. Injections lasted 120 sec with a final off time of 20 minutes at 50 uL/min.

RBD binding to antibodies was assessed using a Series S CM5 chip (Cytiva, MA) which was labeled with anti-human IgG (fc) antibody using a Human Antibody Capture Kit (Cytiva, MA). IgG’s were coated at 200 nM (120s at 5 µL/min) on the chip surface. For single injection experiments, RBDs were injected at 200 nM over the antibodies using the high-performance injection mode (60 sec on, 120 sec off, 50 uL/min). The surface was regenerated three pulses of a 3 M MgCl2 solution for 10 seconds at 100µL/min. Kinetics experiments were performed using single cycle kinetics mode with five different concentrations of analyte serially diluted two-fold. WT RBD concentrations ranged from 0.5 – 8 nM, while XBB RBDs ranged from 2 – 32 nM. Injections lasted 60 sec with a final off time of 120 seconds at 50 uL/min.

Fab binding to Spike proteins was assessed using a Series S SA chip (Cytiva, MA). Spike were coated on the surface at 200 nM (120s at 5 µL/min. For single injection experiments, Fabs were injected at 32 nM over the chip surface (60s at 50 uL/min) with an off time of 120s. The surface was regenerated three pulses of a 3 M MgCl2 solution for 10 seconds at 100µL/min. Kinetics experiments were performed using single cycle kinetics mode with five different concentrations of each Fab serially diluted two-fold. Fab concentrations ranged from 4 – 64 nM. Injections lasted 60 sec with a final off time of 120 seconds at 50 uL/min.

Sensogram data were analyzed using the BiaEvaluation software (Cytiva, MA).

### Cryo-EM

A QuantiFoil Cu300 R1.2/1.3 grid (Electron Microscopy Sciences, PA) was glow discharged at 15 mA for 15 s with 10 s hold time using a PELCO easiGlow™ Glow Discharge Cleaning System. A 2.5-3.5 µl drop of each sample (1.5-2.0 mg/ml in 2 mM Tris, pH 8.0, 200 mM NaCl, 0.02% NaN_3_, 0.5% glycerol) was applied on the grid. One piece of filter paper was utilized for blotting (2.5 s) after 30 s incubation at 22 °C and 95% humidity. The blotting and plunge freeze in liquid ethane were completed by a Leica EM GP2 plunge freezer (Leica Microsystems). The grids were kept in liquid nitrogen and transferred to Titan Krios (Thermo Fisher) for data collection. CryoSPARC (Punjani et al., 2017) was used for data processing and map construction. ChimeraX, Coot, Phenix were used for model building and refinement (Pettersen et al., 2021, Emsley et al., 2010, Adams et al., 2010).

### Vector Based Structure Analysis

The vector analysis was done as previously described (Henderson et al., 2020), which is based on the Visual Molecular Dynamics (VMD) (Humphrey et al., 1996) software package Tcl interface. Briefly, in each protomer, the domain centroids of S1, S2 connector domain, a β-sheet in S2 were determined by Cα, which were then used to calculate the vector distances, angles, and dihedrals. Center and scale of data, principal components analysis, K-means clustering, and Pearson correlation (confidence interval 0.95, p < 0.05) analysis of vectors sets were performed in R (Team, 2018).

### Difference distance matrices (DDM)

DDM were generated using the Bio3D package (Grant et al., 2020) implemented in R (R Core Team (2014). R: A language and environment for statistical computing. R Foundation for Statistical Computing, Vienna, Austria. URL http://www.R-project.org/)

## QUANTIFICATION AND STATISTICAL ANALYSIS

Cryo-EM data were processed and analyzed using cryoSPARC. Cryo-EM structural statistics were analyzed with Phenix and Molprobity. Statistical details of experiments are described in method details or Figure Legends.

