## Supplemental Files for "SARS-CoV-2 Omicron XBB lineage spike structures, conformations, antigenicity, and receptor recognition"

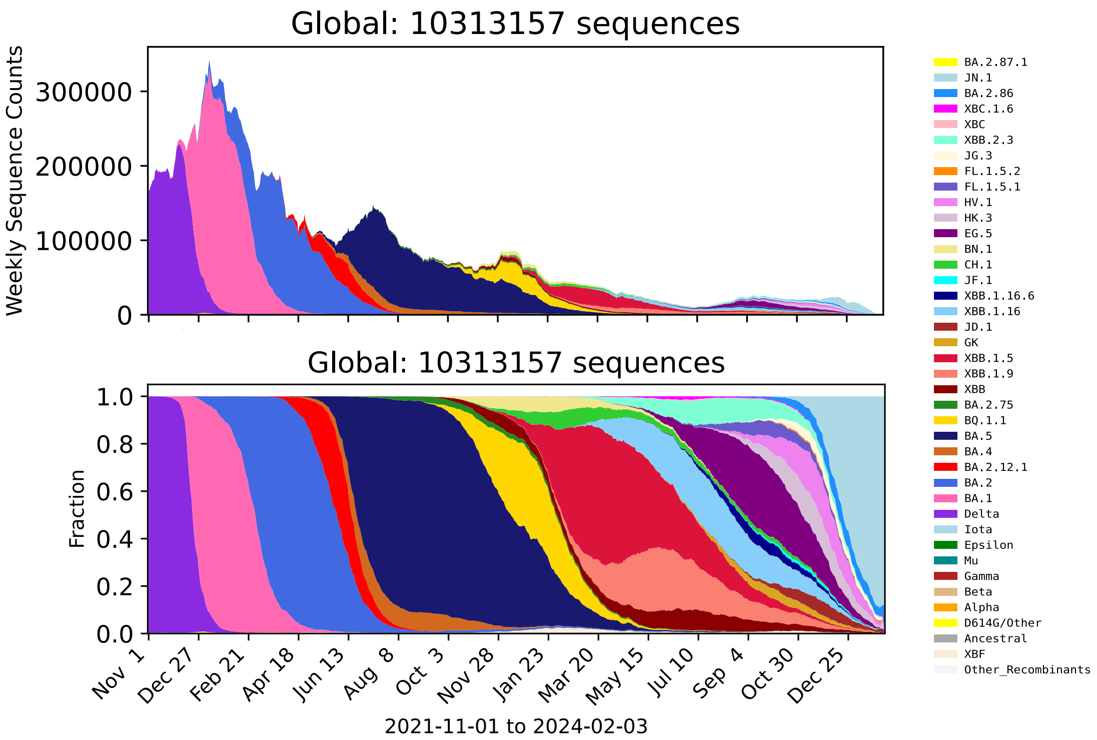
Global: 10313157 sequences
Weekly Sequence Counts
300000
200000
100000
0
Global: 10313157 sequences
Fraction
1.0
0.8
0.6
0.4
0.2
0.0
Nov 1
Dec 27
Feb 21
Apr 18
Jun 13
Aug 8
Oct 3
Nov 28
Jan 23
Mar 20
May 15
Jul 10
Sep 4
Oct 30
Dec 25
2021-11-01 to 2024-02-03
BA.2.87.1
JN.1
BA.2.86
XBC.1.6
XBC
XBB.2.3
JG.3
FL.1.5.2
FL.1.5.1
HV.1
HK.3
EG.5
BN.1
CH.1
JF.1
XBB.1.16.6
XBB.1.16
JD.1
GK
XBB.1.5
XBB.1.9
XBB
BA.2.75
BQ.1.1
BA.5
BA.4
BA.2.12.1
BA.2
BA.1
Delta
Iota
Epsilon
Mu
Gamma
Beta
Alpha
D614G/Other
Ancestral
XBF
Other_Recombinants

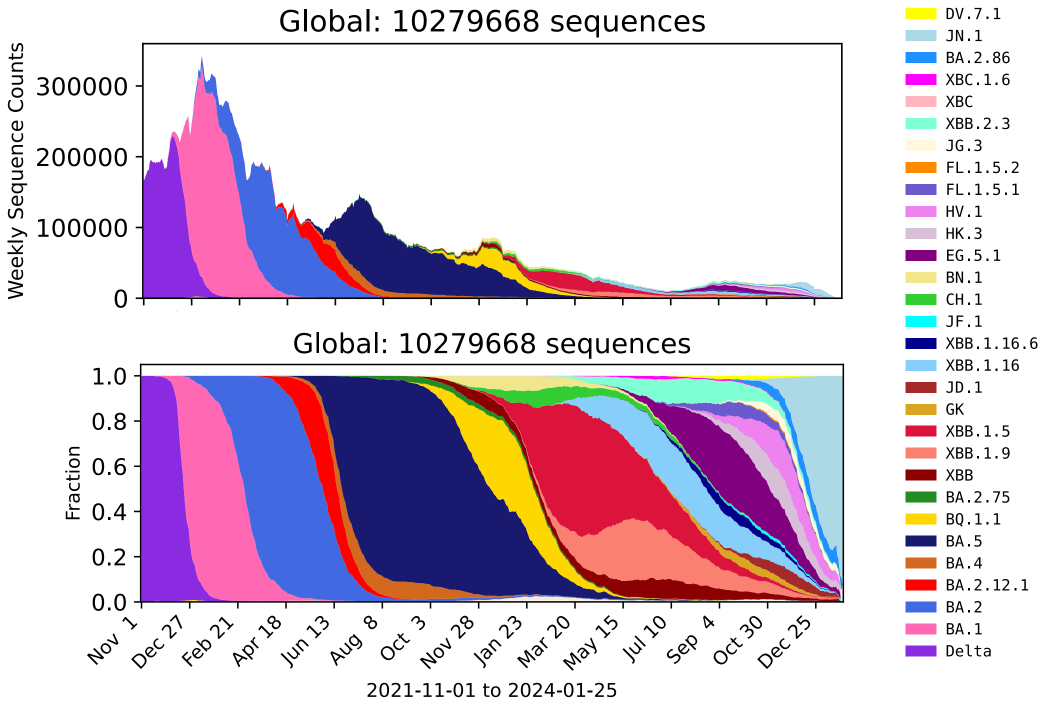
Global: 10279668 sequences
Weekly Sequence Counts
300000
200000
100000
0
Global: 10279668 sequences
Fraction
1.0
0.8
0.6
0.4
0.2
0.0
Nov 1
Dec 27
Feb 21
Apr 18
Jun 13
Aug 8
Oct 3
Nov 28
Jan 23
Mar 20
May 15
Jul 10
Sep 4
Oct 30
Dec 25
2021-11-01 to 2024-01-25
DV.7.1
JN.1
BA.2.86
XBC.1.6
XBC
XBB.2.3
JG.3
FL.1.5.2
FL.1.5.1
HV.1
HK.3
EG.5.1
BN.1
CH.1
JF.1
XBB.1.16.6
XBB.1.16
JD.1
GK
XBB.1.5
XBB.1.9
XBB
BA.2.75
BQ.1.1
BA.5
BA.4
BA.2.12.1
BA.2
BA.1
Delta

**Figure S1: SARS-CoV-2 evolution and global spread.** Global transitions between different Omicron lineages between Nov. 1, 2021, and Jan. 25, 2024, based on 10,564,327 sequences available in this time-period through GISAID (www.gisaid.org). The figure was created using the Embers tool at the COVID-19 Viral Genome Analysis Pipeline (https://cov.lanl.gov/). The presented lineages that are represented here include all sublineages within a given parental lineage, excluding internal sublineages which were expanding and so are singled out and given their own color. The top graphic shows the average weekly counts of sequences, the bottom their frequencies.

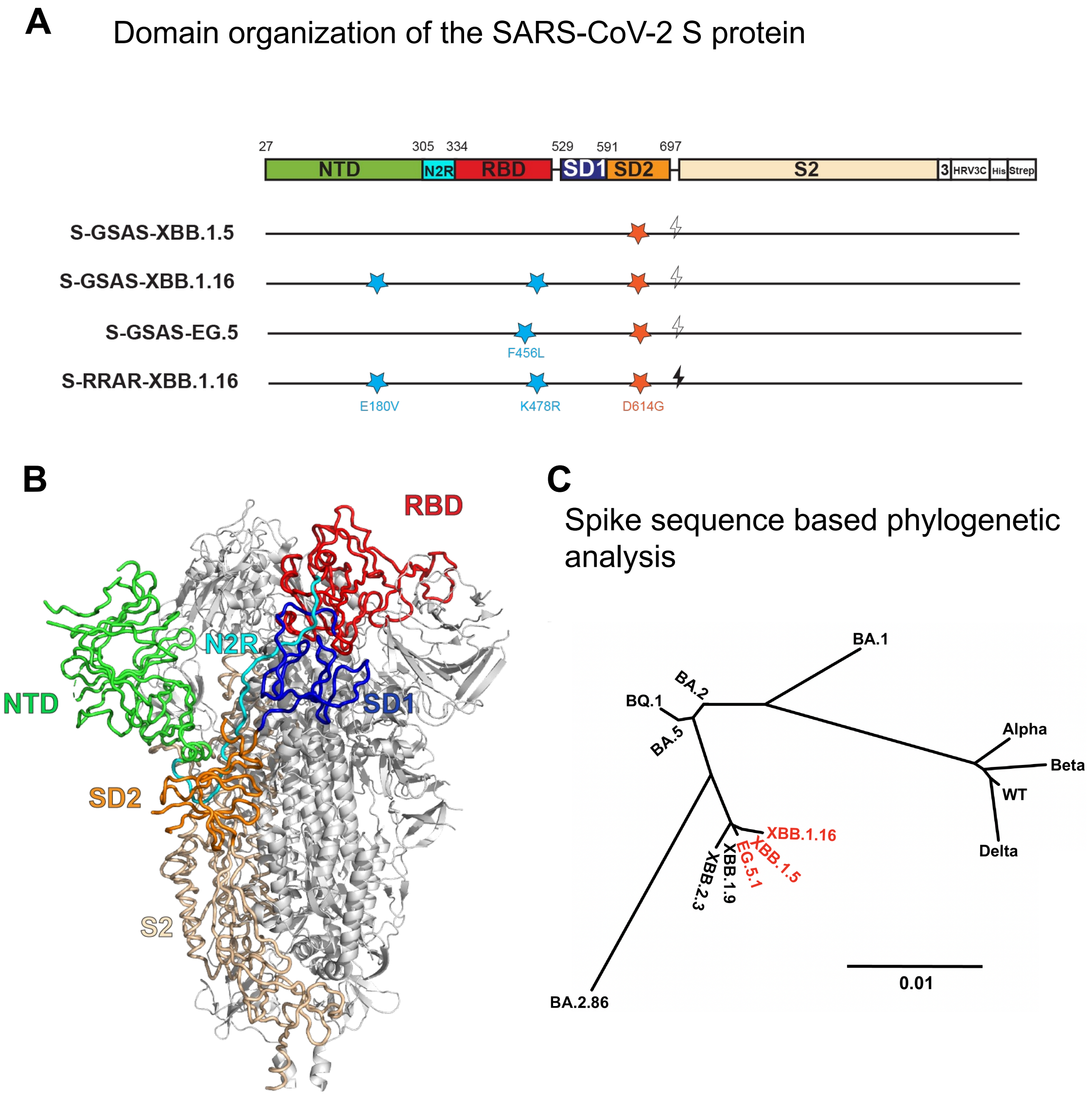


**Figure S2: SARS-CoV-2 S protein domain organization, A.** Domain Organization of SARS-CoV-2 S protein. The S1 subunit contains the NTD (N-terminal domain, pale green), N2R (NTD-to-RBD linker, cyan), RBD (receptor-binding domain, red), and SD1 and SD2 (subdomains 1 and 2, dark blue and orange) subdomains. The S2 subunit is colored wheat. The transmembrane domain (TM) and cytoplasmic tail (CT) have been truncated and replaced by a foldon trimerization sequence, an HRV3C cleavage site (HRV3C), a his-tag (His), and a strep-tag (Strep). The D614G mutation is in the SD2 domain (red star). The S1/S2 furin cleavage site (RRAR; black lightning) has been mutated to GSAS (clear lightning). **B.** S protein structure colored by domains as indicated in panel **A. C.** Phylogenetic tree based of S protein consensus sequences.

**A**

| | DSF | DSC | |
|---|---|---|---|
| | Inflection Temperature (℃) | Melting Temperature (℃) | ΔH (KJ/mol) |
| **WT** | 54.3 ± 0.1 | 51.52 ± 0.01 | 410.8 ± 1.4 |
| **BA.1** | 47.6 ± 0.2 | ND | |
| **BA.2** | 50.6 ± 0.1 | ND | |
| **BA.5** | 49.8 ± 0.2 | ND | |
| **XBB.1.5** | 55.84 ± 0.1 | 53.51 ± 0.02 | 442.6 ± 2.6 |
| **XBB.1.16** | 55.82 ± 0.1 | 53.19 ± 0.02 | 439.3 ± 3.2 |
| **EG.5** | 52.50 ± 0.1 | 50.80 ± 0.01 | 447.5 ± 2.2 |

**B**

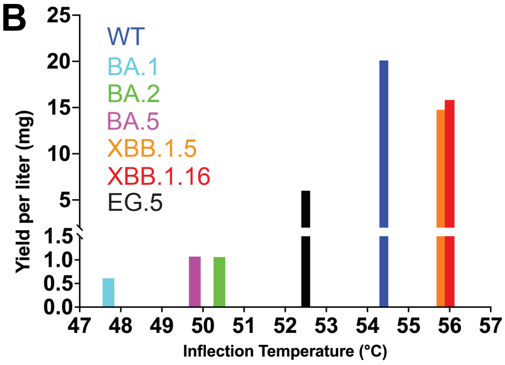


**C**

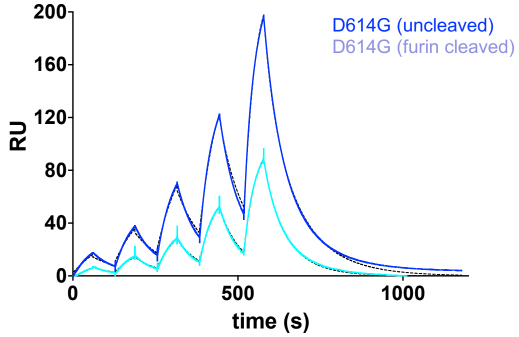


| | ACE2 600s off | | |
|---|---|---|---|
| | ka ($M^{-1}s^{-1}$) | kd ($s^{-1}$) | KD (nM) |
| **D614G (uncleaved)** | 7.27E+05 | 0.08928 | 1.23E-07 |
| **D614G (cleaved)** | 1.77E+05 | 0.03516 | 1.98E-07 |

**D**

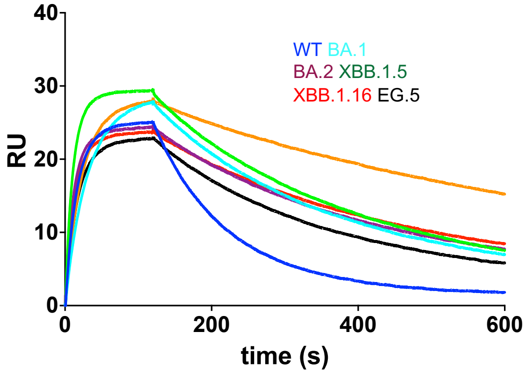


**E**

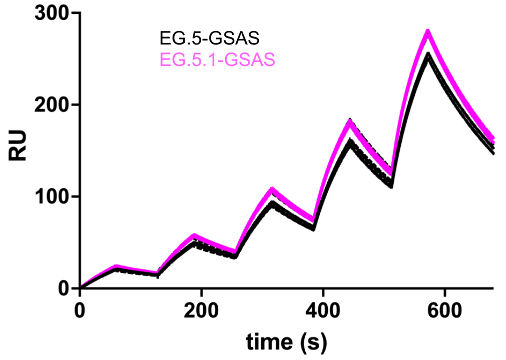


| | ACE2 600s off | | |
|---|---|---|---|
| | ka ($M^{-1}s^{-1}$) | kd ($s^{-1}$) | KD (nM) |
| **EG.5-GSAS** | 6.94E+04 | 0.004924 | 7.10E-08 |
| **EG.5.1-GSAS** | 8.31E+04 | 0.005155 | 6.21E-08 |

**Figure S3: RBD thermostability and ACE2 binding. A.** Table of infection temperatures (Ti) measured by DSF and melting temperature (TM) and enthalpy change of unfolding (delH) measured by DSC in Figure 1C. **B.** Yield of RBD constructs plotted against their thermostability (Ti measured by DSF). **C.** ACE2 binding to uncleaved and furin-cleaved SARS-CoV-2 D614G S ectodomain measured by SPR using the single cycle kinetics format. Experiment was performed exactly as the experiment shown in Figure 2F only with an extended dissociation time of 600 seconds instead of 120 seconds. The raw sensorgrams are shown in dark blue and light blue for the uncleaved and cleaved S ectodomain, respectively. Kinetic parameters obtained from fitting the raw sensorgrams to a 1:1 Langmuir binding model (fitted curves are shown as dotted black lines overlaid on the raw sensorgrams) are shown below the sensograms. **D.** Binding of RBD to ACE2 measured by SPR. ACE2 was amine-coupled to a CM5 sensor chip and RBD at 100 nM concentration were injected over the chip for 2 minutes followed by a 20-minute dissociation phase. **E.** ACE2 binding to EG.5 and EG.5.1 S ectodomain measured by SPR using the same setting as **Figure 2F**.

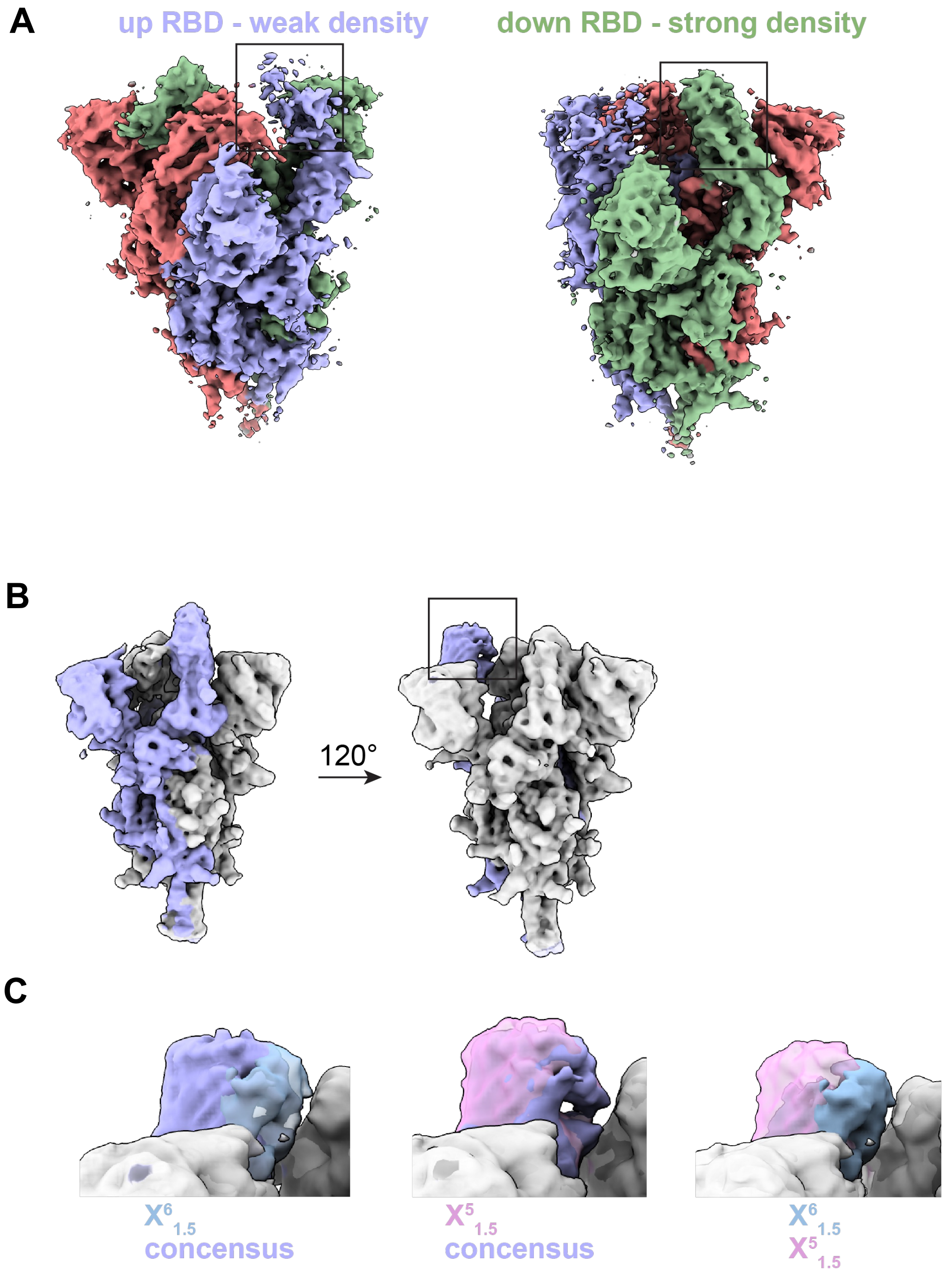


**Figure S4. Variability in the 1-RBD-up state of the XBB.1.5 S ectodomain. A.** Cryo-EM reconstruction of the consensus 1-RBD-up population of XBB.1.5. The up-RBD protomer is colored blue. The two down-RBD protomers are colored red and green. **B.** Gaussian map filter applied to the XBB.1.5 1-RBD-up consensus reconstruction. The down-RBD protomers are colored gray. **C.** Zoomed-in views showing overlay of (left) the consensus and the $X^6_{1.5}$ reconstructions, (middle) the consensus and the $X^5_{1.5}$ reconstructions, (right) the $X^5_{1.5}$ and the $X^6_{1.5}$ reconstructions.

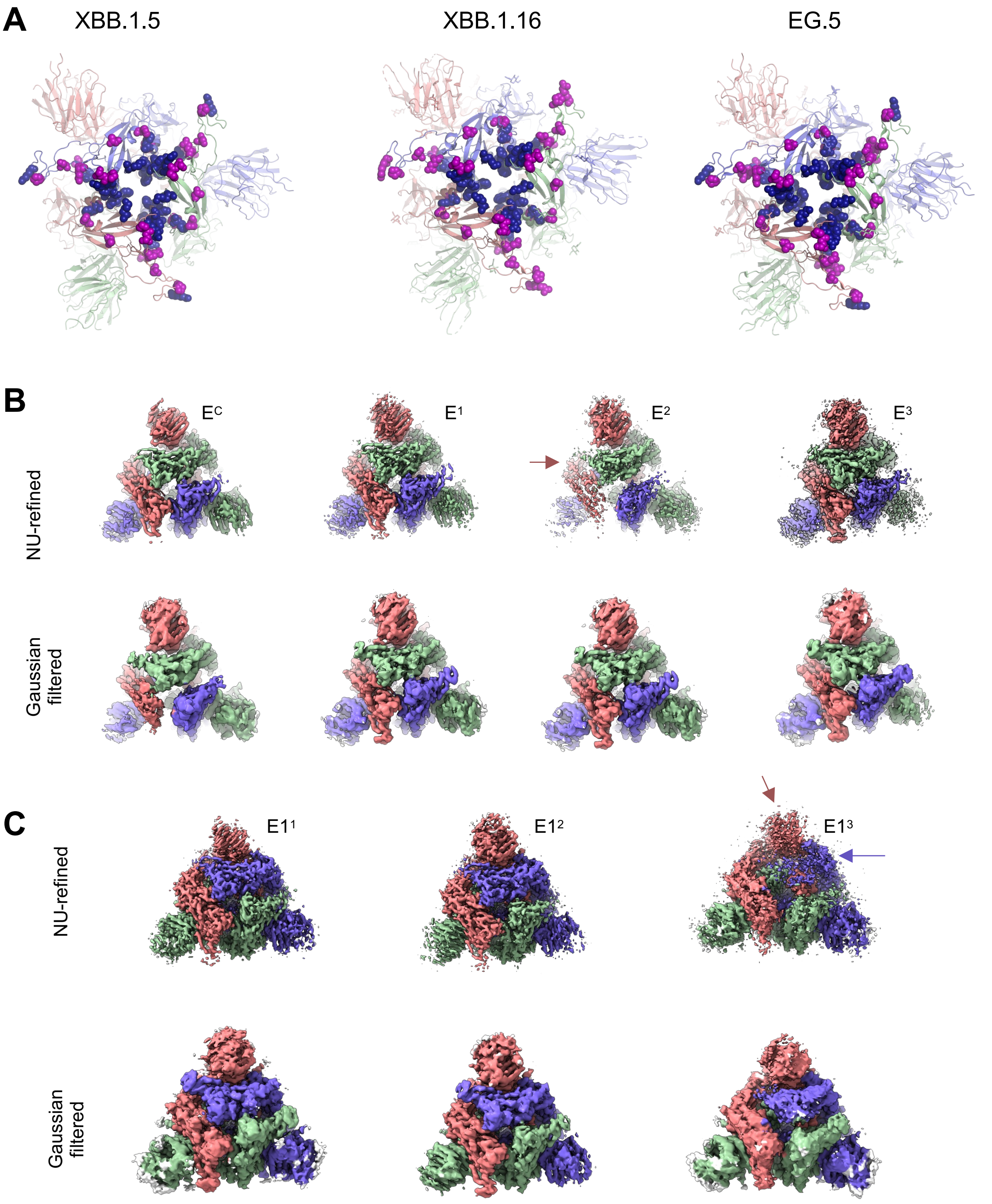


**Figure S5. 3-RBD-down interprotomer packing in XBB.1.5, XBB.1.16, EG.5 and EG.5.1 S proteins.** **A.** 3-RBD-down structures shown for the XBB.1.5 (PDB: 8V0R), XBB.1.16 (PDB:8V0U) and EG.5 (PDB: 8V0V) S ectodomains. RBD mutations are shown as spheres. Mutations common between each structure and BA.2 are colored dark blue. Mutations distinct from BA.2 are colored magenta. **B.** Top views of the 3-RBD-down reconstructions of the EG.5 S ectodomain shown as (top) the non uniform (NU) refinement maps and (bottom) gaussian filtered maps. For $E^2$, the most disordered RBD is indicated by an arrow. **C.** Top views of the 3-RBD-down reconstructions of the EG.5.1 S ectodomain shown as (top) the non uniform (NU) refinement maps and (bottom) gaussian filtered maps. For $E1^3$, the most disordered RBD and NTD are indicated by arrows.

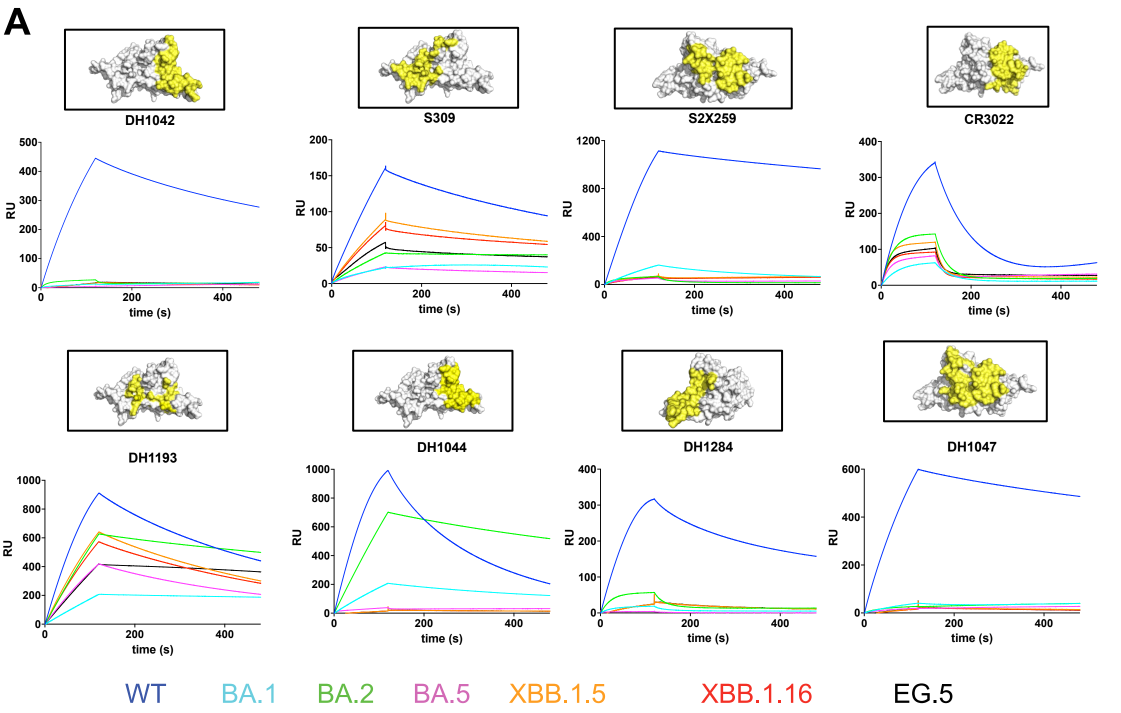


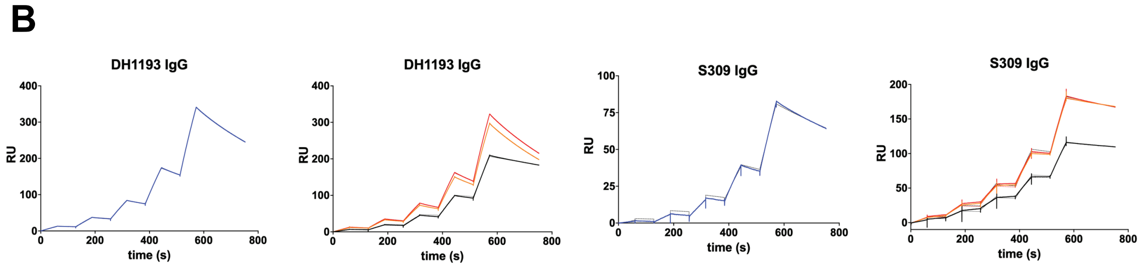


| | WT RBD | | | XBB.1.5 RBD | | | XBB.1.16 RBD | | | EG.5 RBD | | |
|---|---|---|---|---|---|---|---|---|---|---|---|---|
| | ka ($M^{-1}s^{-1}$) | kd ($s^{-1}$) | KD (nM) | ka ($M^{-1}s^{-1}$) | kd ($s^{-1}$) | KD (nM) | ka ($M^{-1}s^{-1}$) | kd ($s^{-1}$) | KD (nM) | ka ($M^{-1}s^{-1}$) | kd ($s^{-1}$) | KD (nM) |
| DH1193 | 1.61E+06 | 0.006848 | 4.25E-09 | 7.17E+05 | 0.006511 | 9.08E-09 | 1.83E+05 | 0.003817 | 2.08E-08 | 5.83E+05 | 0.04463 | 7.66E-08 |
| S309 | 2.66E+05 | 0.058 | 2.18E-07 | 2.04E+05 | 3.86E-04 | 1.89E-09 | 2.34E+05 | 5.03E-04 | 2.15E-09 | 2.56E+05 | 2.95E-04 | 1.15E-09 |

**Figure S6. Antigenicity of RBD variants. A.** SPR analysis of RBD antigenicity. 200nM of each RBD was flowed over antibodies immobilized on anti-Fc surface. Epitope of each RBD-reactive antibody is mapped in yellow to the RBD surface colored in grey. **B.** Single cycle kinetics sensorgrams of RBD (analyte) interaction with antibody IgG (captured on an anti-Fc coated chip). WT RBD was injected at a concentration range 0.5-8 nM, the XBB RBDs were injected at 2-32 nM. The table shows the affinity and kinetics parameters obtained from fitting the sensorgrams to a Langmuir 1:1 binding model.

A

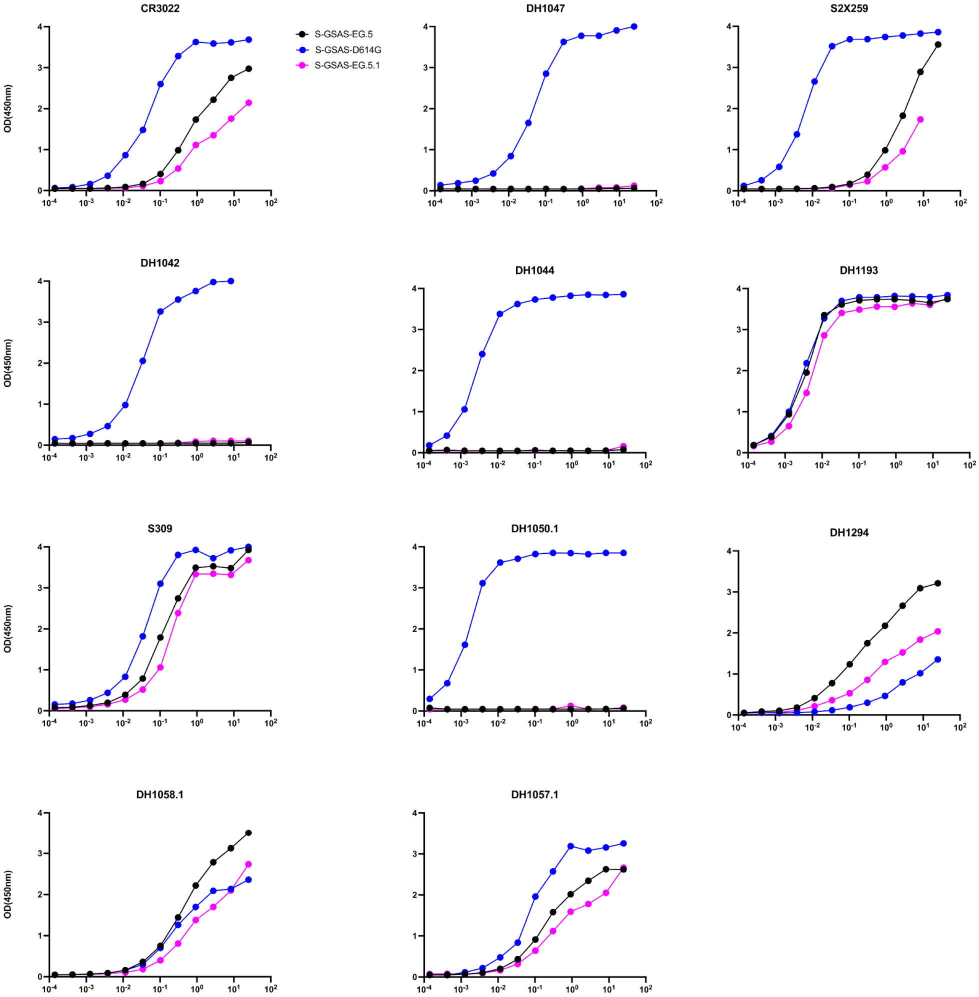


B

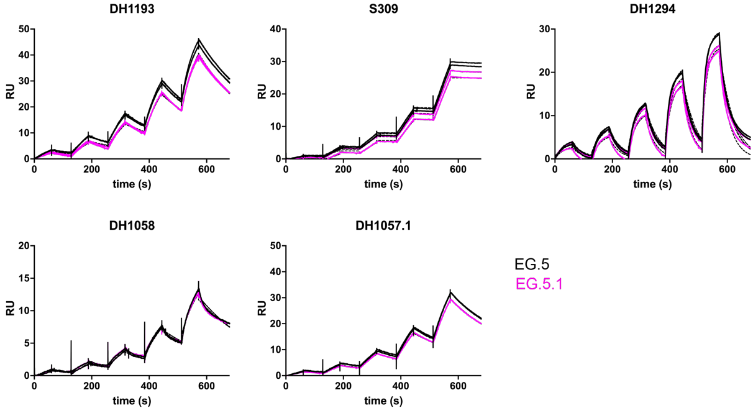


| | EG.5 | | | EG.5.1 | | |
|---|---|---|---|---|---|---|
| | ka ($M^{-1}s^{-1}$) | kd ($s^{-1}$) | KD (nM) | ka ($M^{-1}s^{-1}$) | kd ($s^{-1}$) | KD (nM) |
| **DH1193** | 2.35E+05 | 3.86E-03 | 1.65E-08 | 2.47E+05 | 5.12E-03 | 2.08E-08 |
| **S309** | 4.32E+04 | 1.54E-04 | 3.80E-09 | 2.09E+04 | 7.73E-05 | 2.24E-09 |
| **DH1294** | 8.27E+05 | 2.64E-02 | 3.19E-08 | 7.66E+05 | 3.01E-02 | 3.93E-08 |
| **DH1058.1** | 1.12E+05 | 4.55E-03 | 4.23E-08 | 1.25E+05 | 4.38E-03 | 3.84E-08 |
| **DH1057.1** | 1.11E+05 | 3.57E-03 | 3.26E-08 | 8.41E+04 | 3.78E-03 | 4.50E-08 |

**Figure S7: Comparison of EG.5 and EG.5.1 S ectodomain antigenicity. A.** Binding of uncleaved D614G, EG.5 and EG.5.1 S ectodomains to a panel of antibodies measured by ELISA**. B.** Binding of uncleaved D614G, EG.5 and EG.5.1 S ectodomains to a panel of antibodies measured by SPR using the single cycle kinetics format**.**

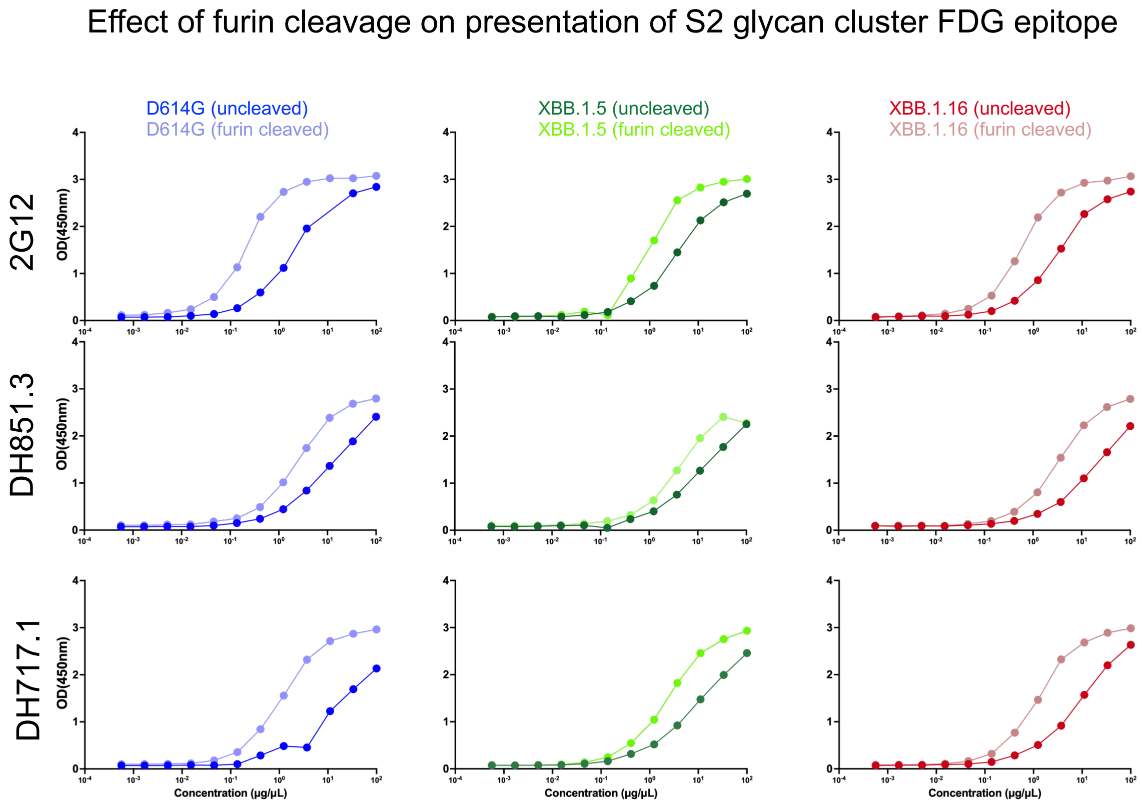


**Figure S8. FDG antibody binding to SARS-CoV-2 S ectodomains.** ELISA binding of FDG antibodies to uncleaved and furin-cleaved S ectodomains.

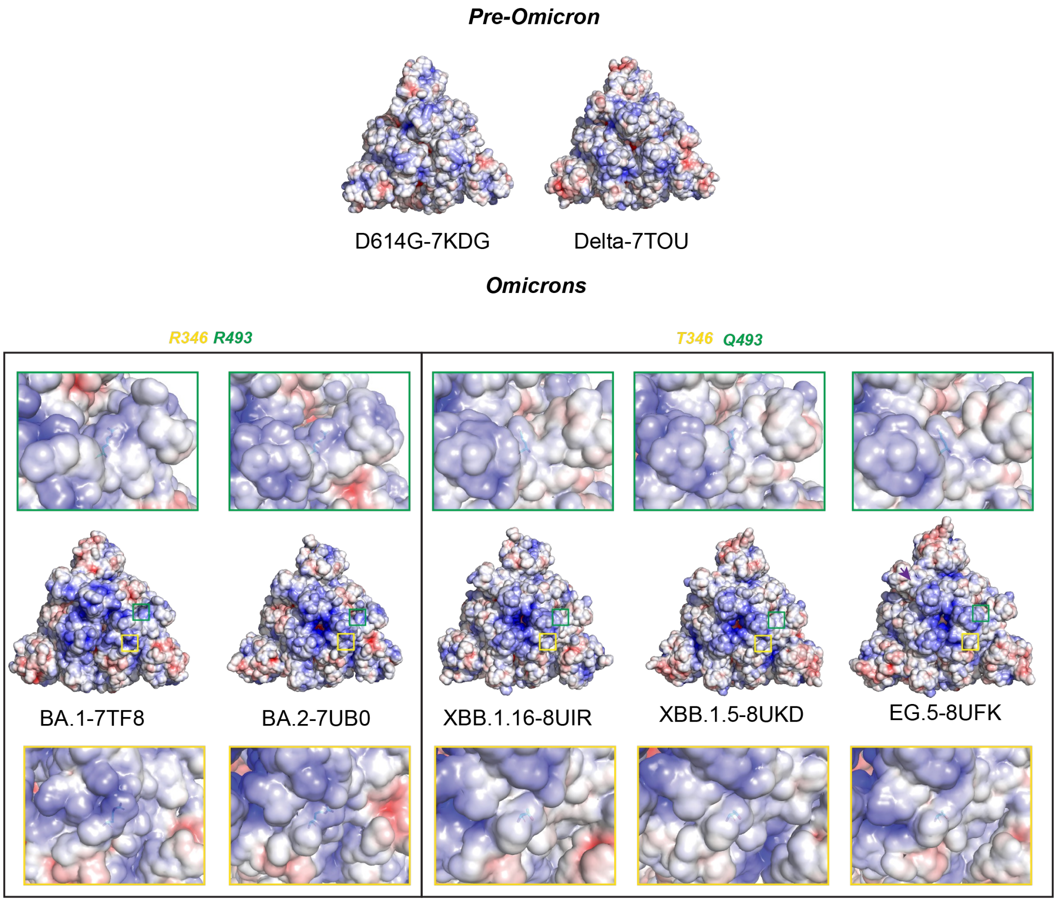


**Figure S9. Evolution of surface electrostatics in Omicron S proteins.** Surface representations of S ectodomains are shown colored by their electrostatic potentials. The top panel shows the D614G and Delta spikes (pre-Omicron) and the bottom panels show Omicron spikes. Top views are shown looking down from the host cell membrane. Yellow and green squares show the locations of the R346T and R493Q substitutions that contributed to decrease in overall surface electropositivity, with a electropositive patch now concentrated at the center of the spike surface.

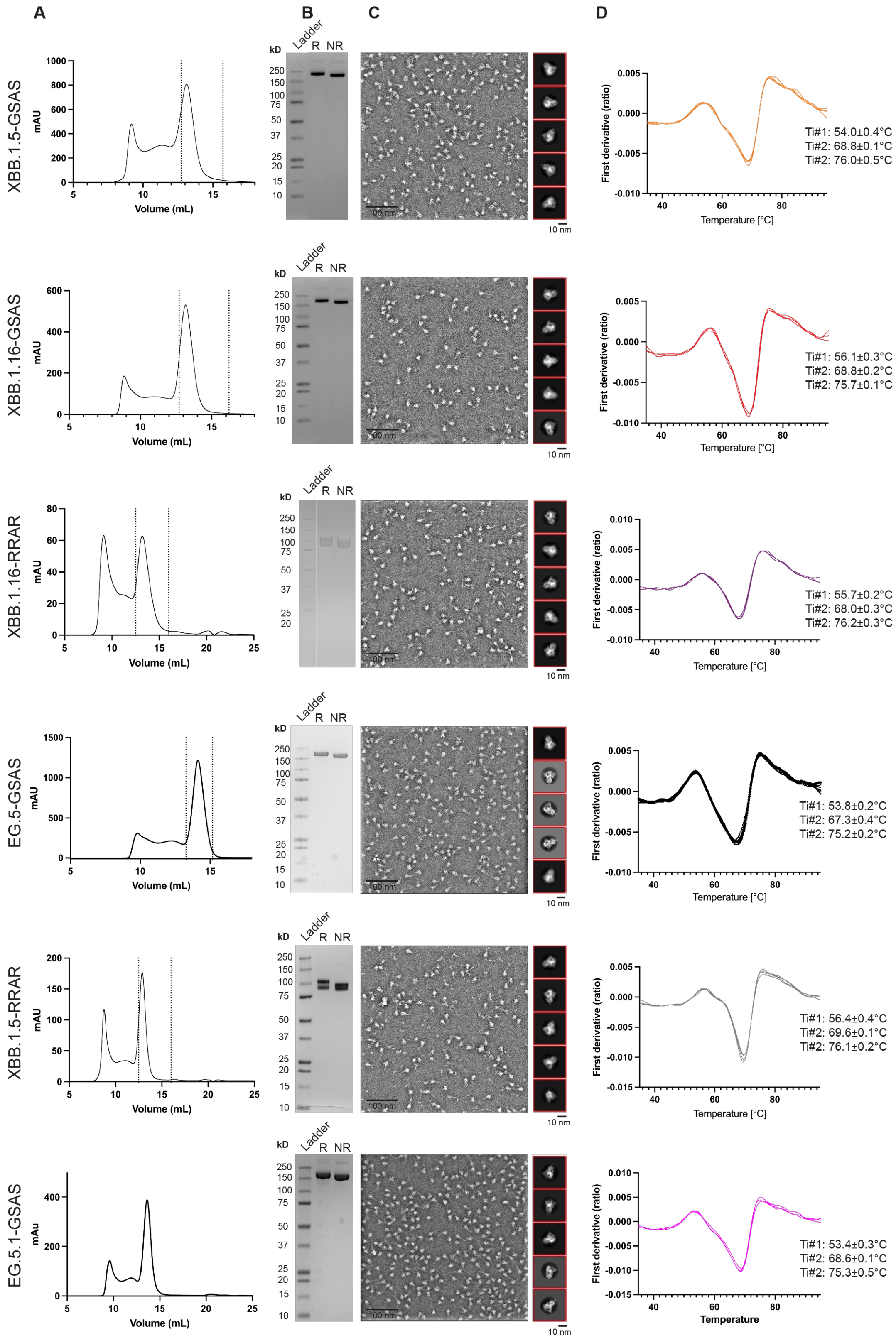
A
B
C
D
XBB.1.5-GSAS
XBB.1.16-GSAS
XBB.1.16-RRAR
EG.5-GSAS
XBB.1.5-RRAR
EG.5.1-GSAS
mAU
Volume (mL)
Ladder
R
NR
kD
250
150
100
75
50
37
25
20
15
10
100 nm
10 nm
First derivative (ratio)
Temperature [°C]
Temperature
Ti#1: 54.0±0.4°C
Ti#2: 68.8±0.1°C
Ti#2: 76.0±0.5°C
Ti#1: 56.1±0.3°C
Ti#2: 68.8±0.2°C
Ti#2: 75.7±0.1°C
Ti#1: 55.7±0.2°C
Ti#2: 68.0±0.3°C
Ti#2: 76.2±0.3°C
Ti#1: 53.8±0.2°C
Ti#2: 67.3±0.4°C
Ti#2: 75.2±0.2°C
Ti#1: 56.4±0.4°C
Ti#2: 69.6±0.1°C
Ti#2: 76.1±0.2°C
Ti#1: 53.4±0.3°C
Ti#2: 68.6±0.1°C
Ti#2: 75.3±0.5°C

**Data S1: Purification of SARS-CoV-2 S ectodomain proteins.** From left to right. Size exclusion chromatography (SEC) profile. Dashed lines indicate the fractions pooled for final products. SDS-PAGE. Lane 1, Molecular weight marker; lane 2, reduced 2 µg of spike; lane 3, non-reduced 2 µg of spike. Negative Stain Electron Micrograph (NSEM) with representative 2D classes shown on the right. Differential scanning fluorimetry (DSF) plots of spikes, with three technical replicates shown for XBB.1.16 and EG.5.1, four for XBB.1.5, and six for EG.5. The average and standard deviation of three temperatures of inflection were calculated.

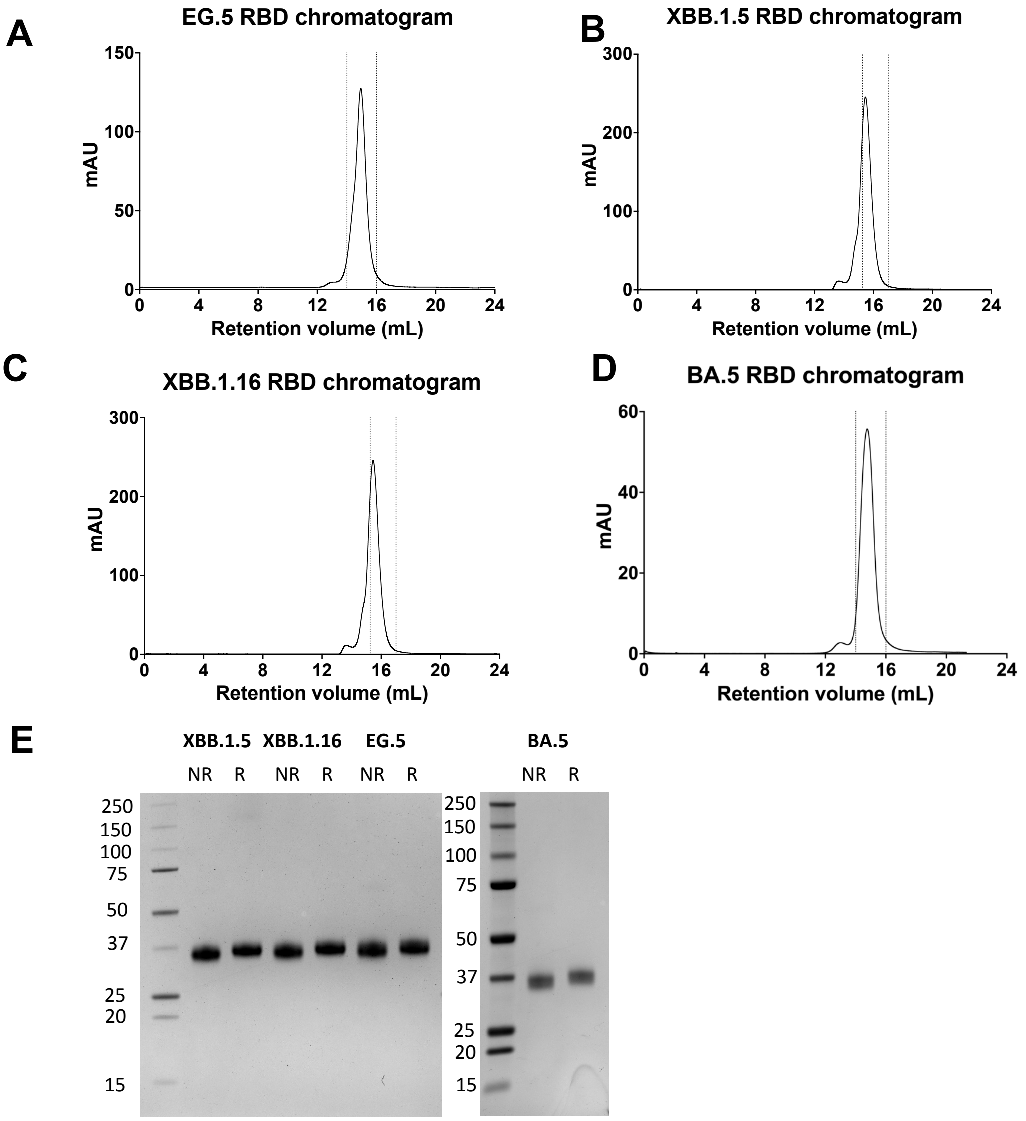


**Data S2. Purification of SARS-CoV-2 S protein Receptor Binding Domains (RBD). A-C.** Size exclusion chromatogram of the RBD of **A.** EG.5 **B.** XBB.1.5 and **C.** XBB.1.16. **D.** BA.5. **E.** SDS-PAGE of purified RBD.

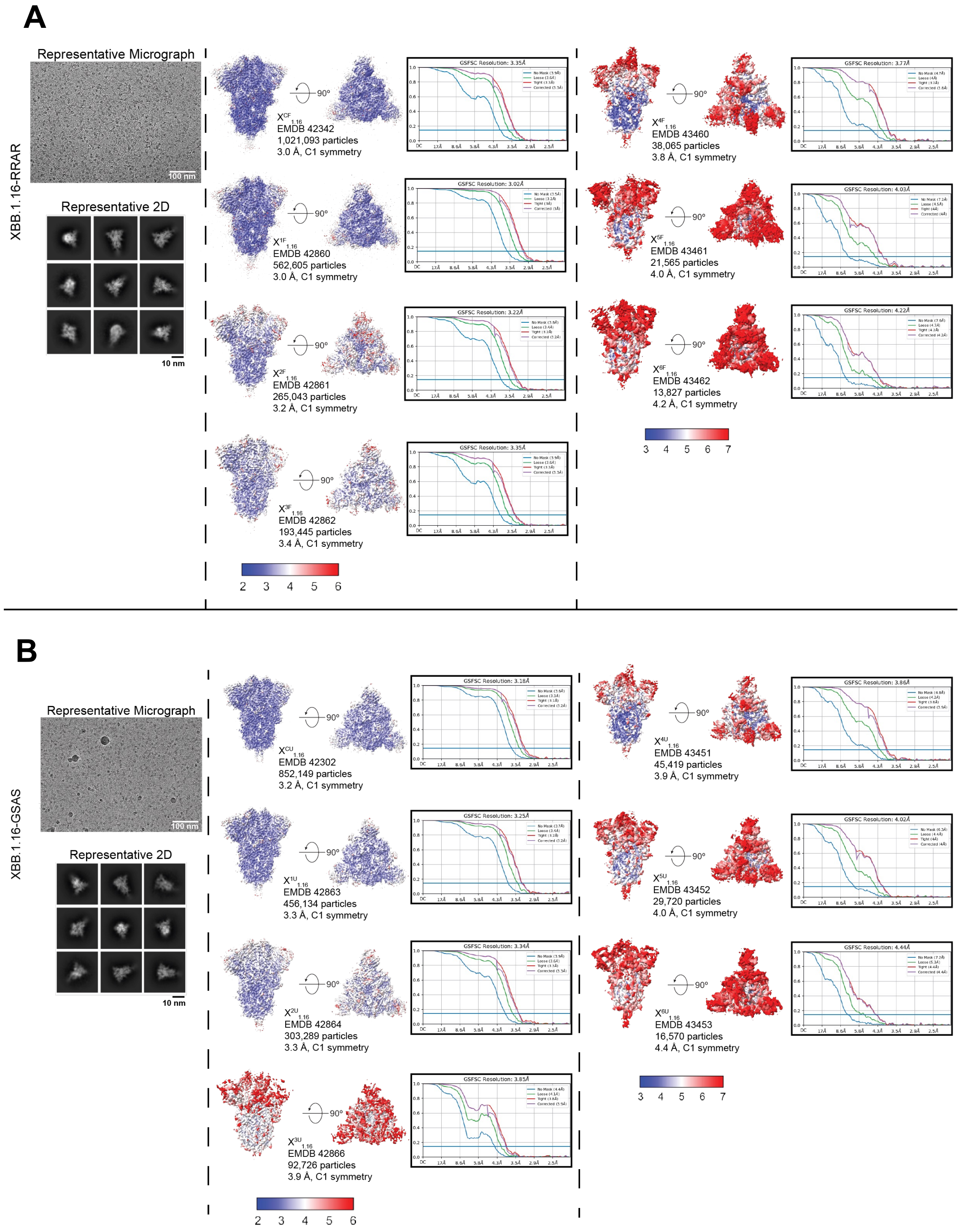


**Data S3. Cryo-EM data processing and map quality of the S ectodomains,** Representative micrographs and 2D class averages (left), local resolution estimation and FSC plot of 3-RBD-down maps (middle), local resolution estimation and FSC plot of 1-RBD-up maps (right) are shown for datasets of (A) XBB.1.16-RRAR, (B) XBB.1.16-GSAS, (C) XBB.1.5-GSAS, (D) EG.5-GSAS, (E) EG.5.1-GSAS. Graphs are generated by cryoSPARC and ChimeraX.

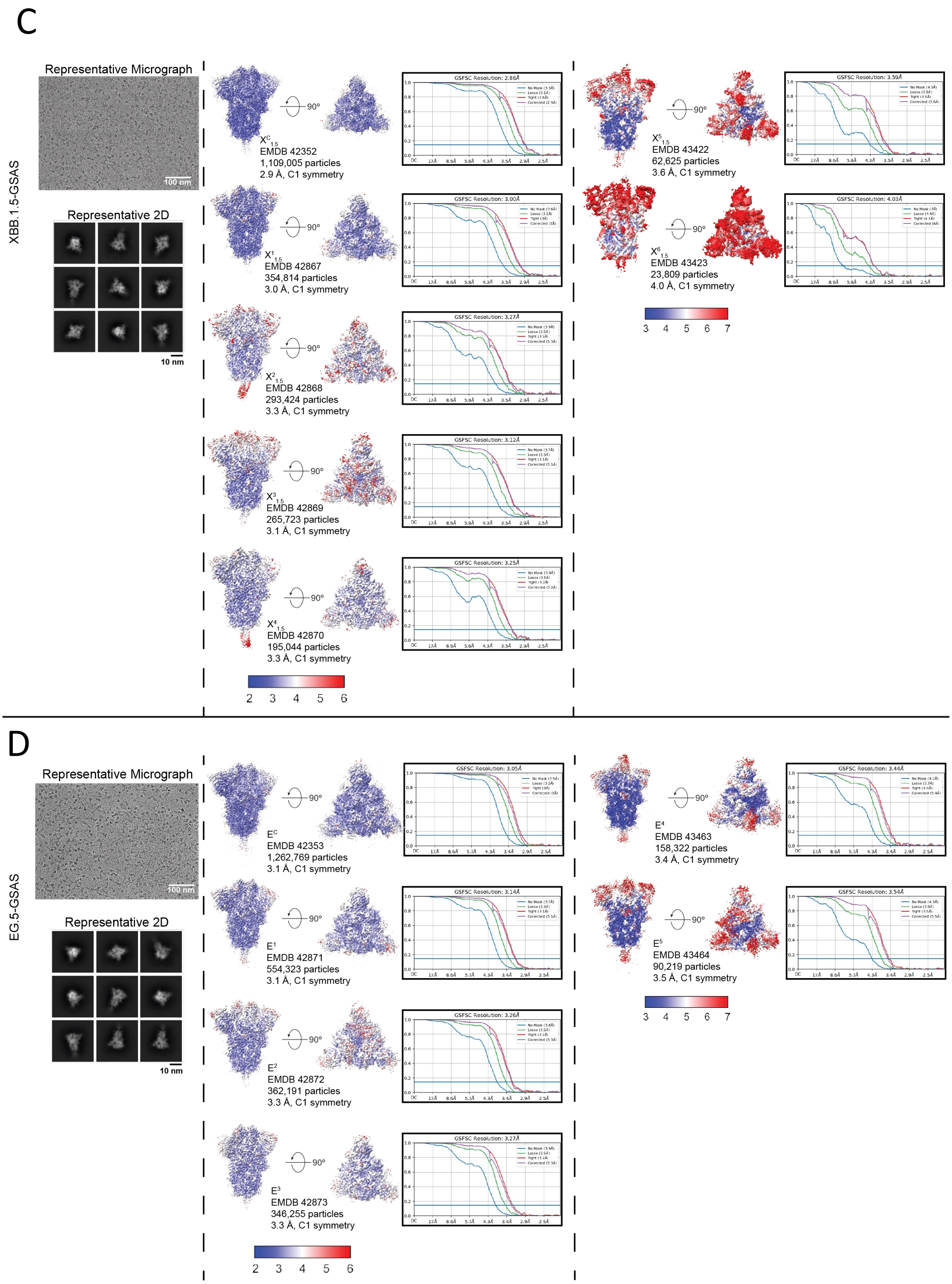


**Data S3 (continued)**

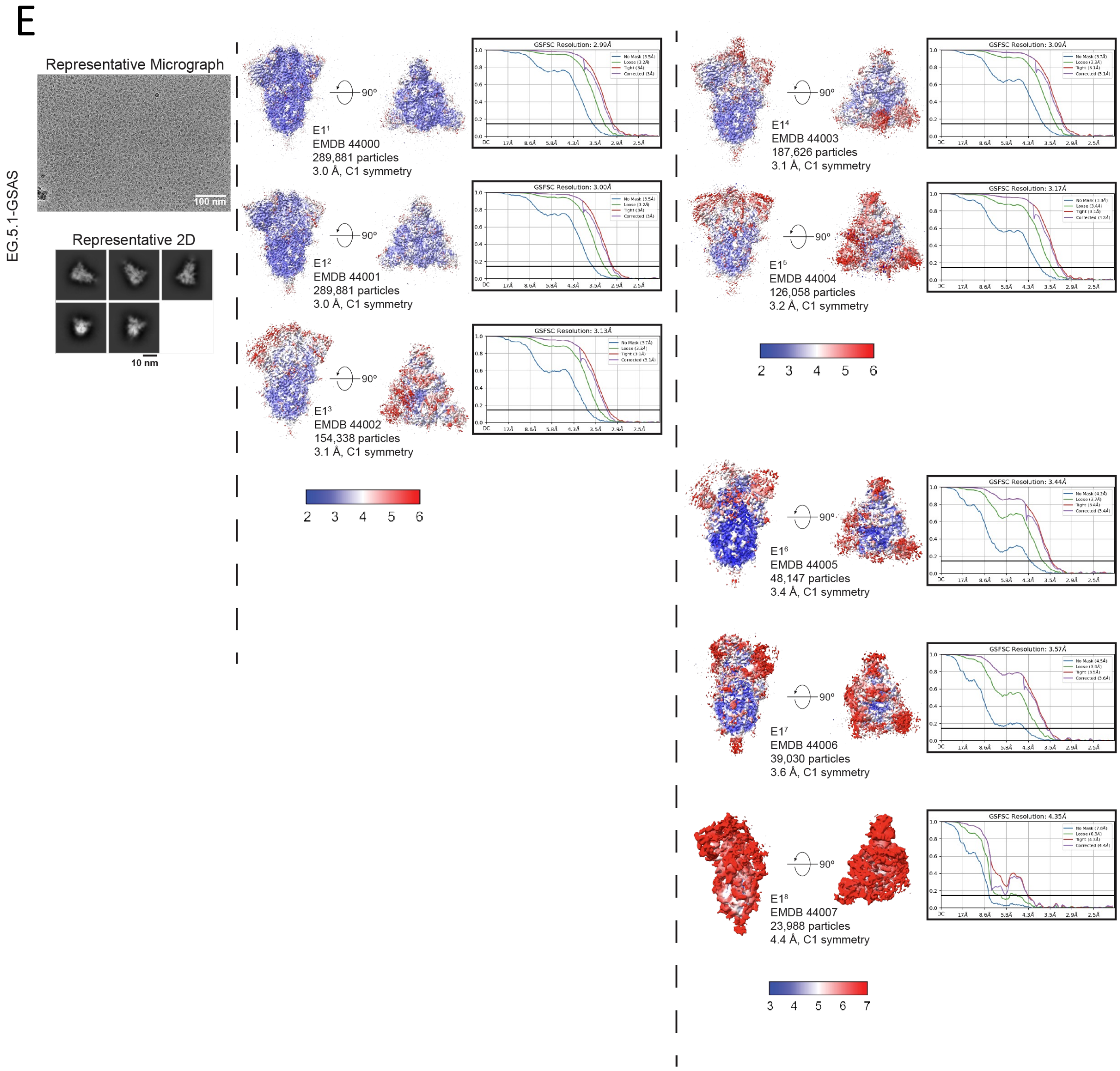


**Data S3 (continued)**

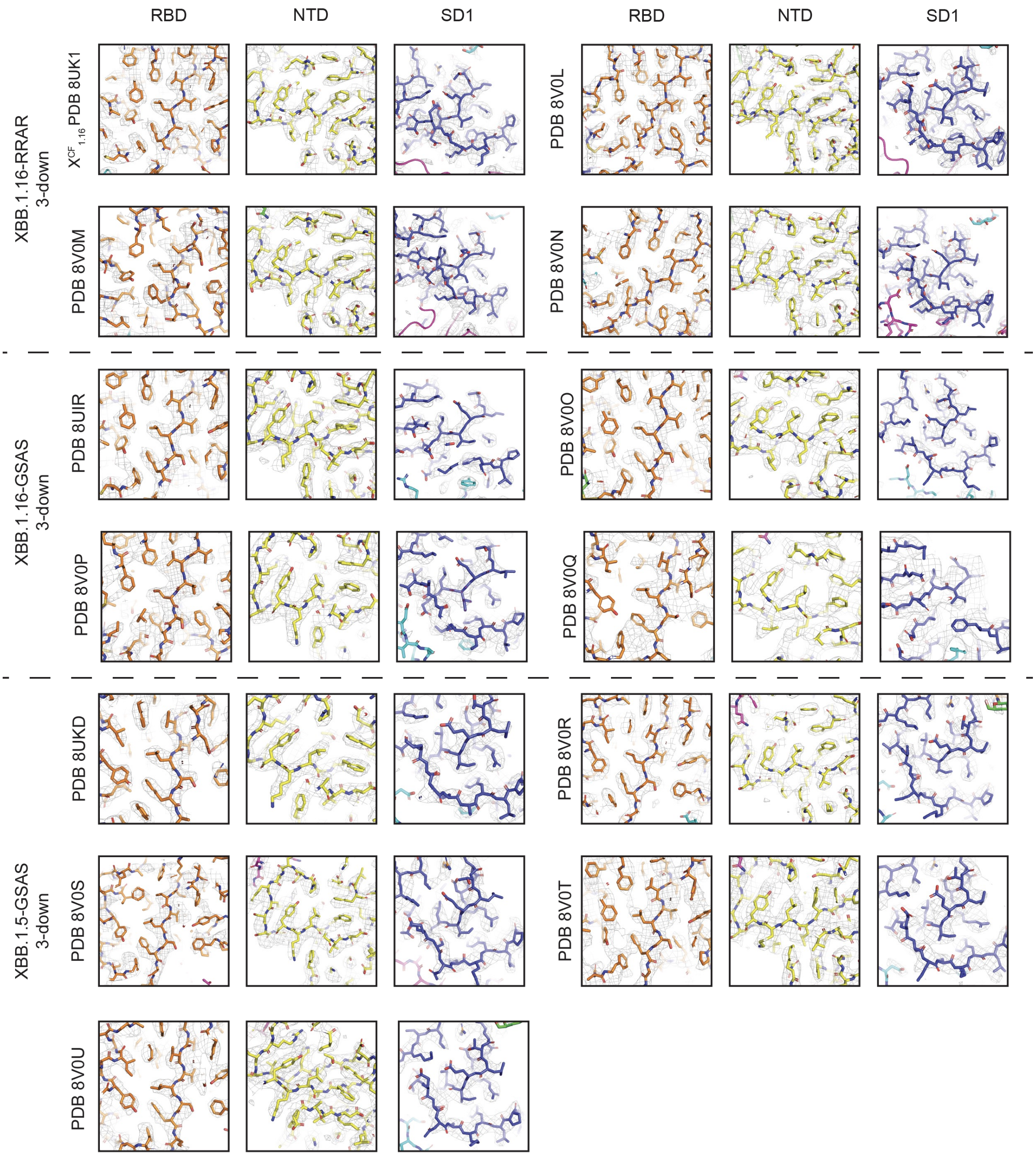
RBD
NTD
SD1
RBD
NTD
SD1
XBB.1.16-RRAR
3-down
$X^{CF}_{1.16}$ PDB 8UK1
PDB 8V0L
PDB 8V0M
PDB 8V0N
XBB.1.16-GSAS
3-down
PDB 8UIR
PDB 8V0O
PDB 8V0P
PDB 8V0Q
PDB 8UKD
PDB 8V0R
XBB.1.5-GSAS
3-down
PDB 8V0S
PDB 8V0T
PDB 8V0U

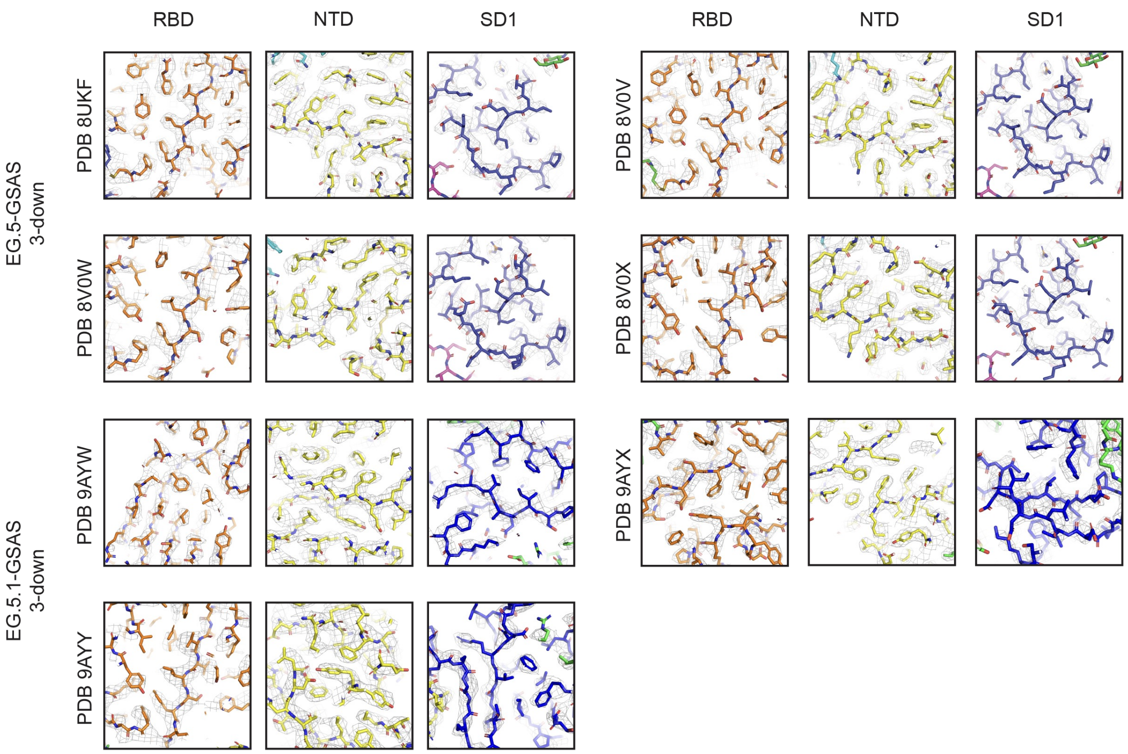


**Data S4. Cryo-EM data quality.** Zoomed in views of different regions of the structure.

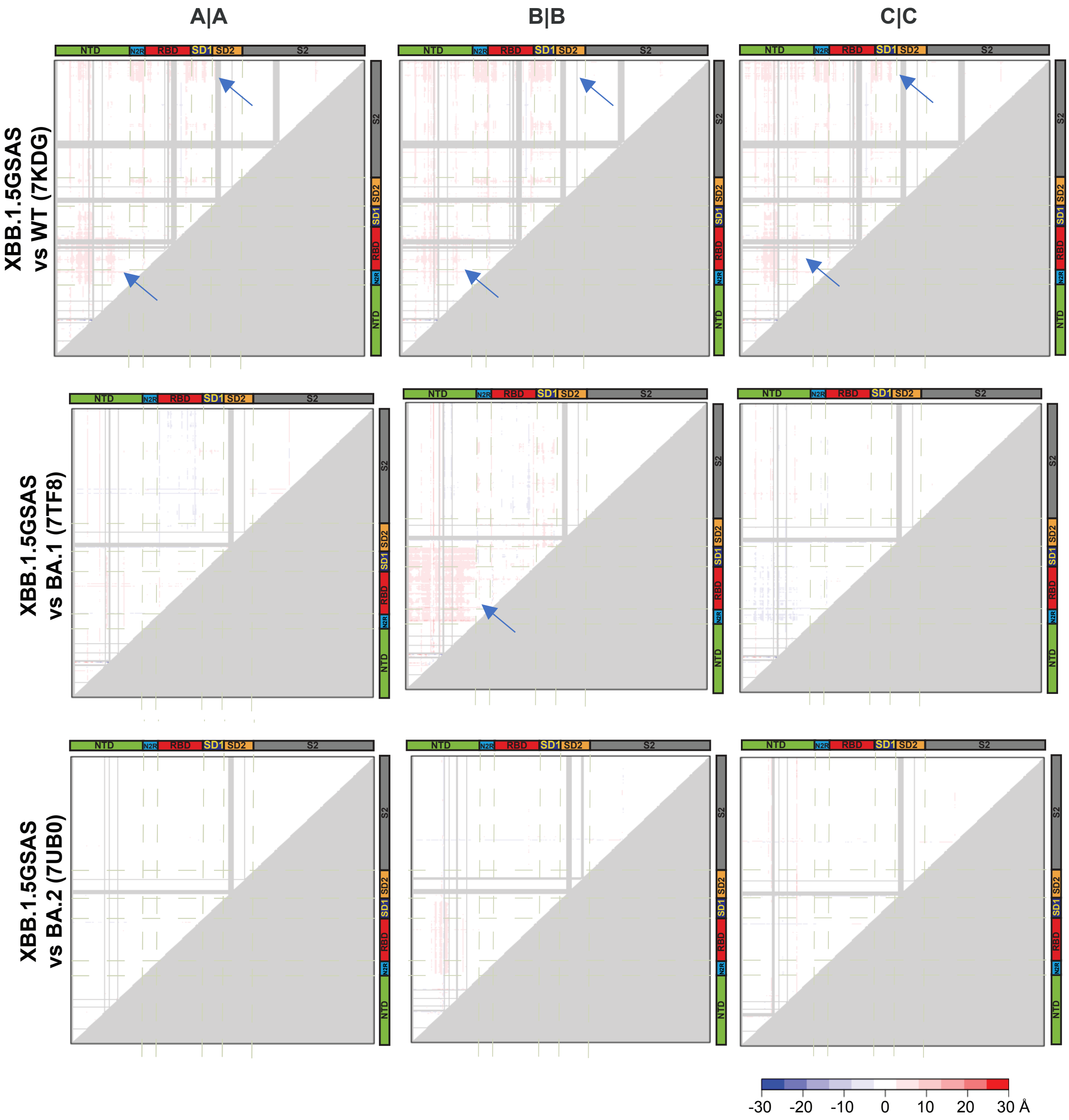


**Data S5. Difference Distance Matrix (DDM) analysis of S-GSAS-XBB.1.5, S-GSAS-XBB.1.16, and S-GSAS-EG.5 compared to previous strains.** The protomers of the consensus 3-RBD-down states in this study were compared with the corresponding protomers of S-GSAS-D614G, S-GSAS-BA.1 and S-GSAS-BA.2. Arrows indicate regions with significant difference distances (> 3Å).

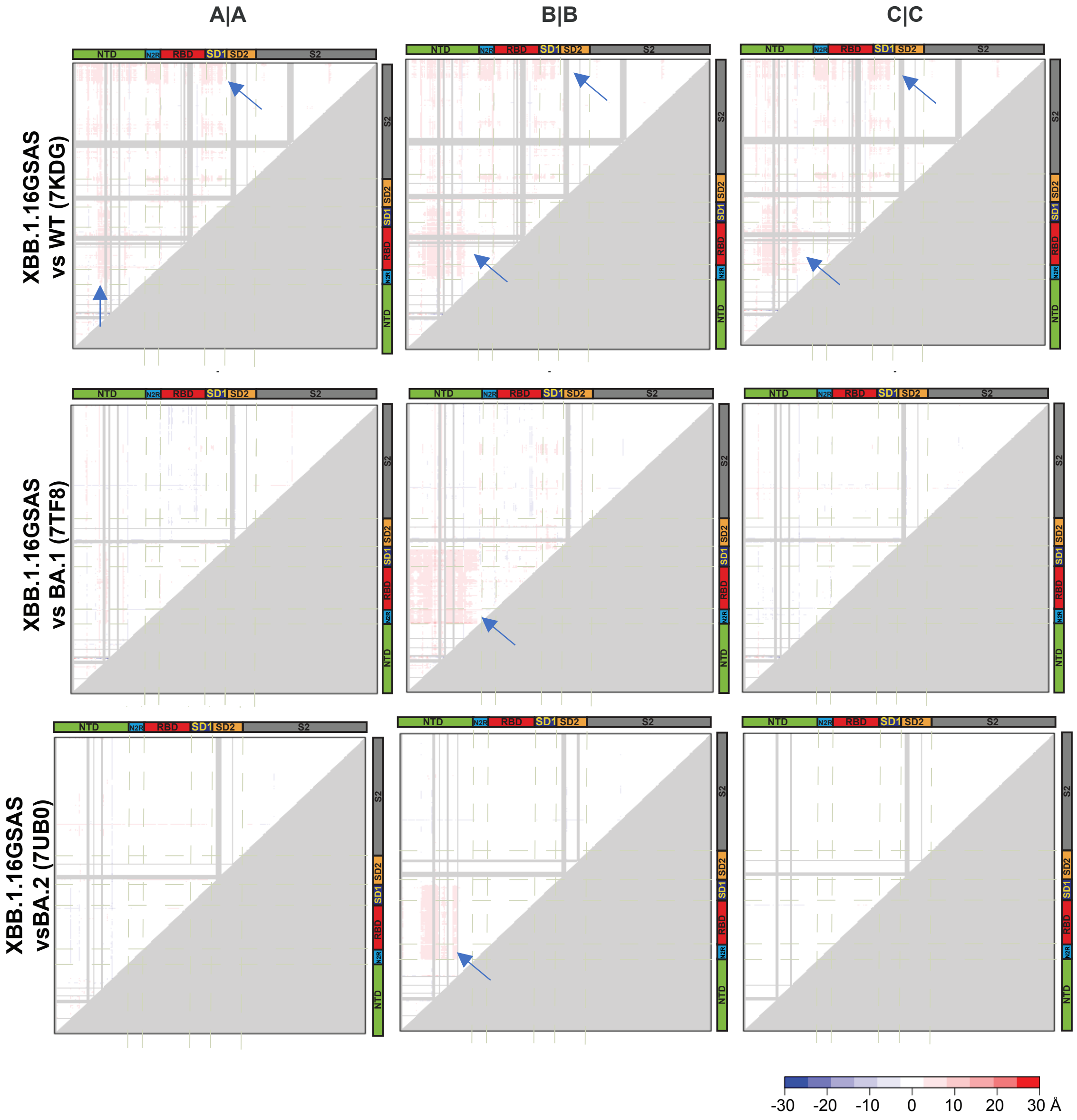


**Data S5 (contd.). Difference Distance Matrix (DDM) analysis of S-GSAS-XBB.1.5, S-GSAS-XBB.1.16, and S-GSAS-EG.5 compared to previous strains.**

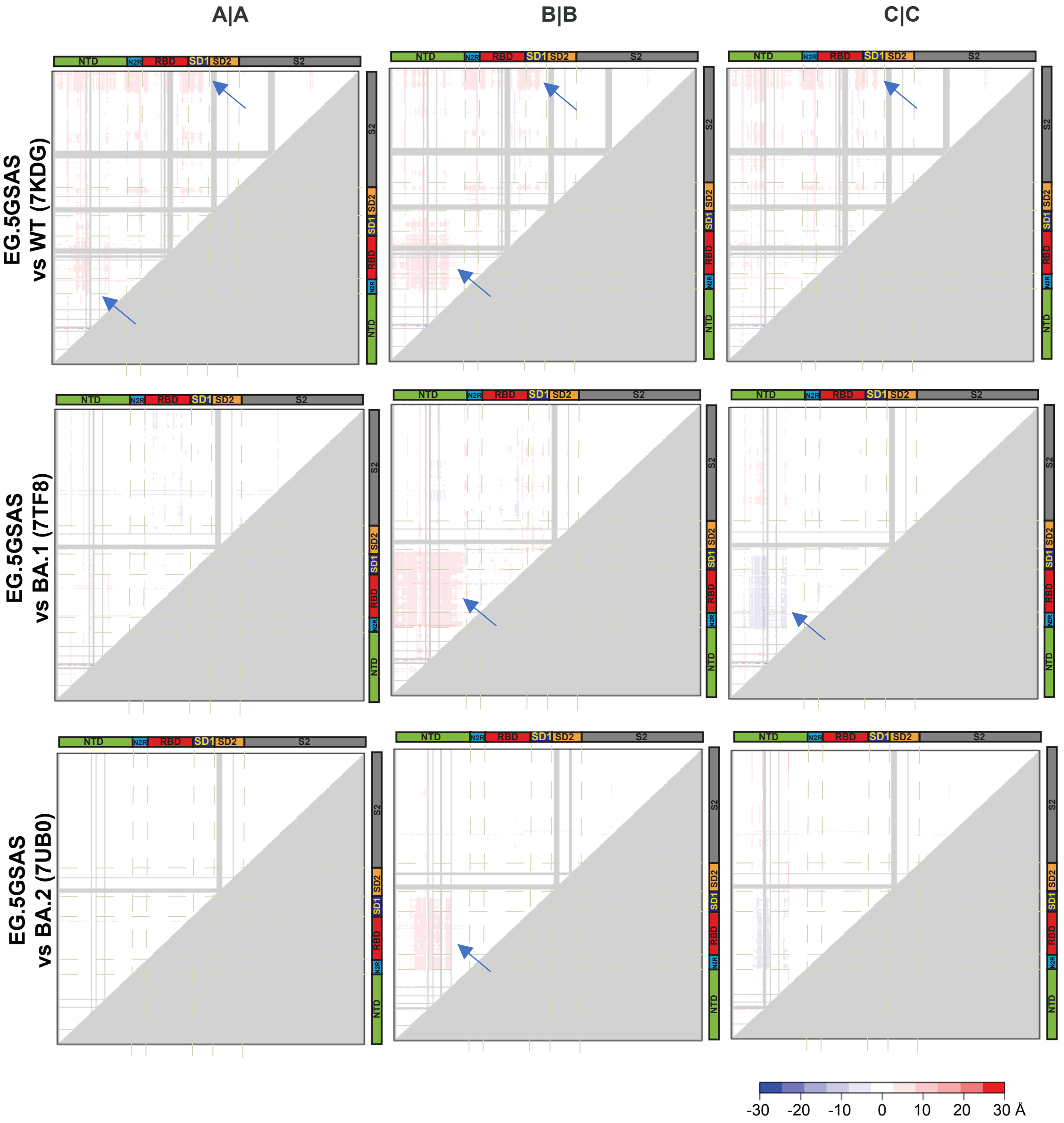


**Data S5 (contd.). Difference Distance Matrix (DDM) analysis of S-GSAS-XBB.1.5, S-GSAS-XBB.1.16, and S-GSAS-EG.5 compared to previous strains.**

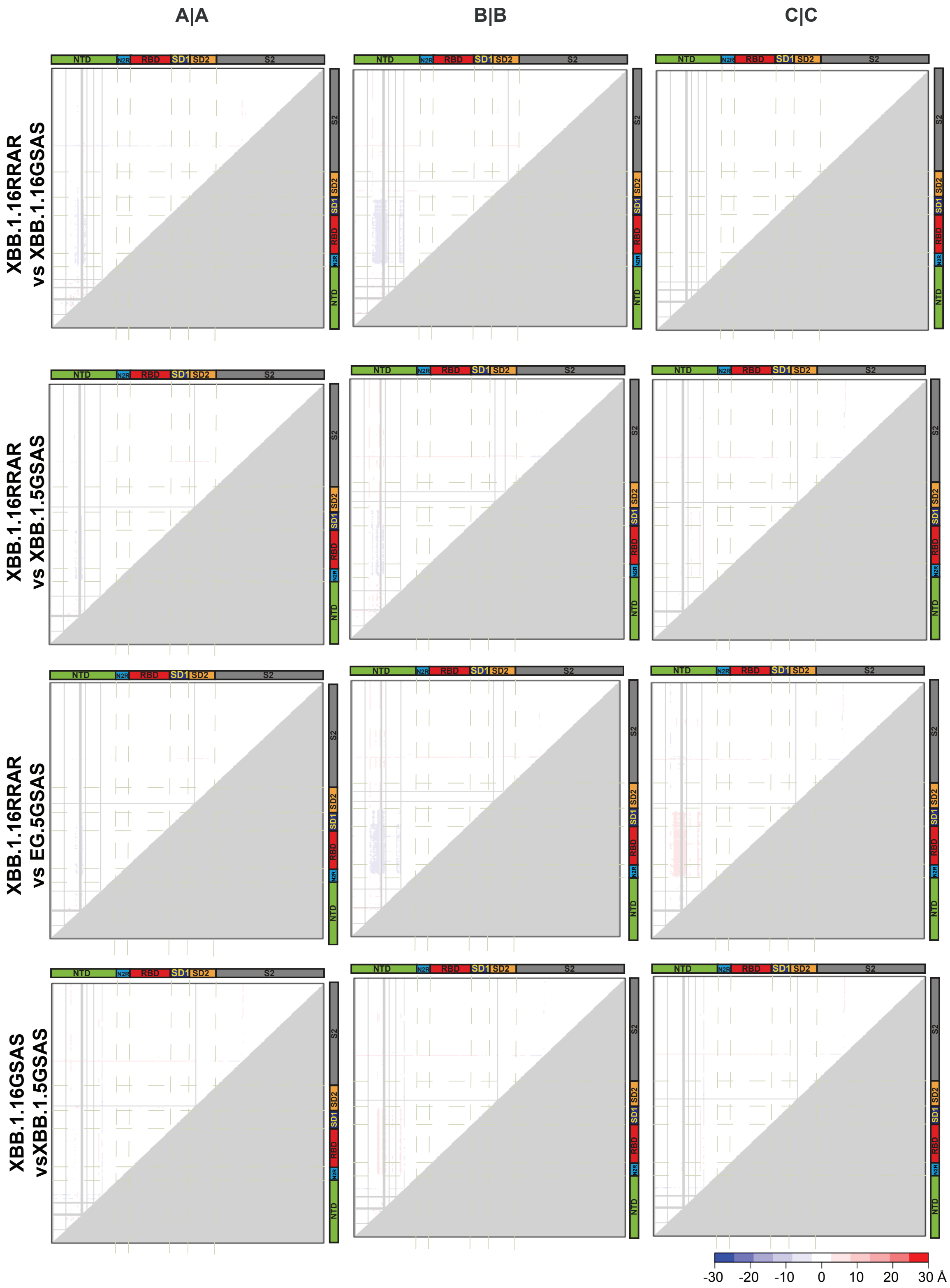


**Data S6. Difference Distance Matrix (DDM) analysis of XBB.1.5, XBB.1.16 and EG.5 spikes.**

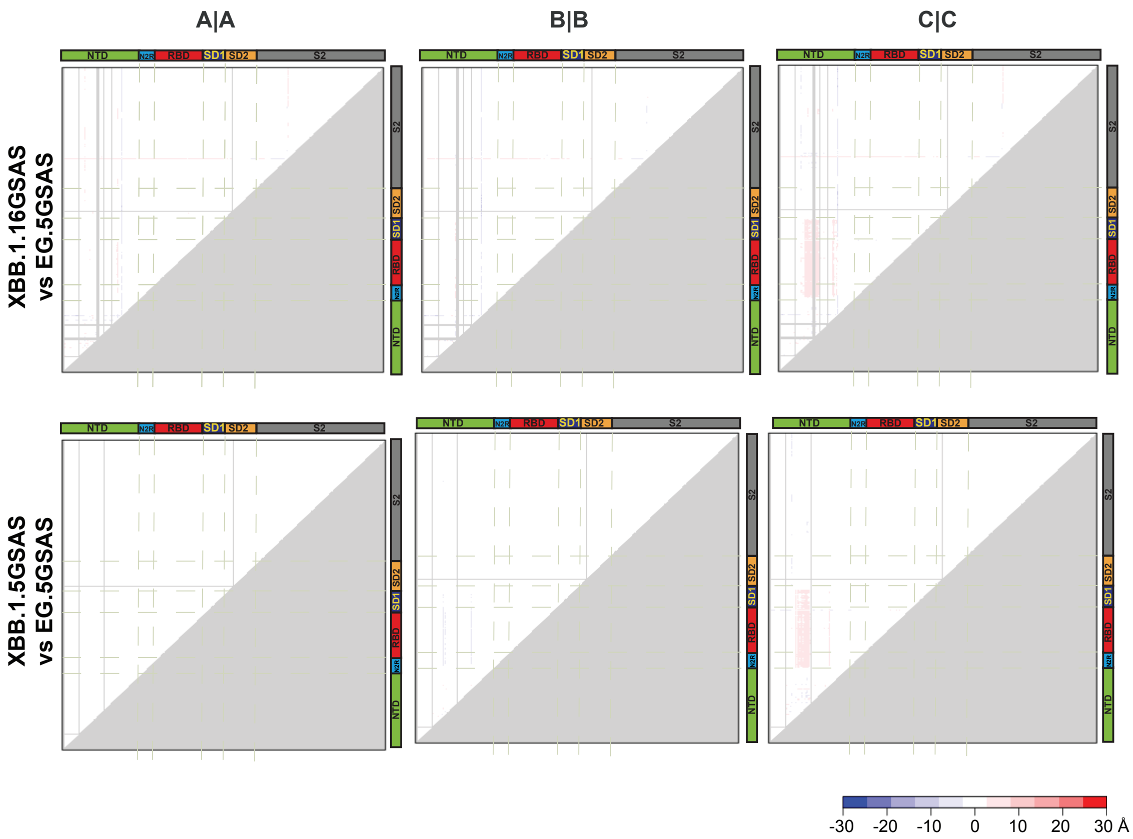


**Data S6 (contd). Difference Distance Matrix (DDM) analysis of XBB.1.5, XBB.1.16 and EG.5 spikes.**

### Table S1: Cryo-EM data collection and refinements statistics

| | S-GSAS/XBB.1.5 | | | | | | | S-GSAS/XBB.1.16 | | | | | | |
|---|---|---|---|---|---|---|---|---|---|---|---|---|---|---|
| | $X^{C}_{1.5}$ | $X^{1}_{1.5}$ | $X^{2}_{1.5}$ | $X^{3}_{1.5}$ | $X^{4}_{1.5}$ | $X^{5}_{1.5}$ | $X^{6}_{1.5}$ | $X^{CU}_{1.16}$ | $X^{1U}_{1.16}$ | $X^{2U}_{1.16}$ | $X^{3U}_{1.16}$ | $X^{4U}_{1.16}$ | $X^{5U}_{1.16}$ | $X^{6U}_{1.16}$ |
| **PDB ID** | **8UKD** | **8V0R** | **8V0S** | **8V0T** | **8V0U** | | | **8UIR** | **8V0O** | **8V0P** | **8V0Q** | | | |
| **EMDB ID** | **42352** | **42867** | **42868** | **42869** | **42870** | **43422** | **43423** | **42302** | **42863** | **42864** | **42866** | **43451** | **43452** | **43453** |
| **Data Collection and processing** | | | | | | | | | | | | | | |
| Microscope | FEI Titan Krios | | | | | | | FEI Titan Krios | | | | | | |
| Detector | Gatan K3 | | | | | | | Gatan K3 | | | | | | |
| Magnification | 81000 | | | | | | | 81000 | | | | | | |
| Voltage (kV) | 300 | | | | | | | 300 | | | | | | |
| Electron exposure (e-/Å^2) | 59 | | | | | | | 59 | | | | | | |
| Defocus Range (µm) | 2.4 to 0.8 | | | | | | | 2.4 to 0.8 | | | | | | |
| Pixel size (Å) | 1.08 | | | | | | | 1.08 | | | | | | |
| Reconstruction software | cryoSPARC | | | | | | | cryoSPARC | | | | | | |
| Micrographs used for data processing | 7,708 | | | | | | | 6,629 | | | | | | |
| Symmetry imposed | C1 | | | | | | | C1 | | | | | | |
| Initial particle images (no.) | 3,638,075 | | | | | | | 2,343,651 | | | | | | |
| Final particle images (no.) | 1,109,005 | 354,814 | 293,424 | 265,723 | 195,044 | 62,625 | 23,809 | 852,149 | 456,134 | 303,289 | 92,726 | 45,419 | 29,720 | 16,570 |
| Map resolution (Å) | 2.9 | 3.0 | 3.3 | 3.1 | 3.3 | 3.6 | 4.0 | 3.2 | 3.3 | 3.3 | 3.9 | 3.8 | 4.0 | 4.4 |
| FSC threshold | | | | | | | | | | | | | | |
| **Model composition** | | | | | | | | | | | | | | |
| Nonhydrogen atoms | 24777 | 24903 | 24950 | 24777 | 24811 | | | 24879 | 24879 | 24925 | 24795 | | | |
| Protein residues | 3123 | 3123 | 3126 | 3123 | 3105 | | | 3105 | 3105 | 3111 | 3105 | | | |
| **R.M.S. deviations** | | | | | | | | | | | | | | |
| Bond lengths (Å) | 0.005 | 0.004 | 0.004 | 0.006 | 0.003 | | | 0.003 | 0.004 | 0.004 | 0.004 | | | |
| Bond angles (°) | 0.818 | 0.719 | 0.863 | 0.807 | 0.734 | | | 0.677 | 0.742 | 0.768 | 0.8 | | | |
| **Validation** | | | | | | | | | | | | | | |
| MolProbity score | 1.39 | 1.36 | 1.8 | 1.51 | 1.44 | | | 1.29 | 1.41 | 1.46 | 1.47 | | | |
| Clashscore | 2.78 | 2.74 | 4.97 | 3.2 | 3.18 | | | 2.38 | 2.93 | 3.21 | 3.47 | | | |
| Rotomer outliers (%) | 0.04 | 0 | 0 | 0.04 | 0 | | | 0.04 | 0 | 0 | 0 | | | |
| **Ramachandran plot** | | | | | | | | | | | | | | |
| Favored regions (%) | 95.29 | 95.68 | 90.53 | 94.13 | 95.23 | | | 96.07 | 95.26 | 95 | 95.29 | | | |
| Ourliers (%) | 0.03 | 0.1 | 0.1 | 0.03 | 0.03 | | | 0 | 0.07 | 0 | 0.03 | | | |

| | S-RRAR/XBB.1.16 | | | | | | | S-GSAS/EG.5 | | | | | |
|---|---|---|---|---|---|---|---|---|---|---|---|---|---|
| | $X^{CF}_{1.16}$ | $X^{1F}_{1.16}$ | $X^{2F}_{1.16}$ | $X^{3F}_{1.16}$ | $X^{4F}_{1.16}$ | $X^{5F}_{1.16}$ | $X^{6F}_{1.16}$ | $E^{C}$ | $E^{1}$ | $E^{2}$ | $E^{3}$ | $E^{4}$ | $E^{5}$ |
| **PDB ID** | **8UK1** | **8V0L** | **8V0M** | **8V0N** | | | | **8UKF** | **8V0V** | **8V0W** | **8V0X** | | |
| **EMDB ID** | **42342** | **42860** | **42861** | **42862** | **43460** | **43461** | **43462** | **42353** | **42871** | **42872** | **42873** | **43463** | **43464** |
| **Data Collection and processing** | | | | | | | | | | | | | |
| Microscope | FEI Titan Krios | | | | | | | FEI Titan Krios | | | | | |
| Detector | Gatan K3 | | | | | | | Gatan K3 | | | | | |
| Magnification | 81000 | | | | | | | 81000 | | | | | |
| Voltage (kV) | 300 | | | | | | | 300 | | | | | |
| Electron exposure (e-/Å^2) | 59 | | | | | | | 72.9 | | | | | |
| Defocus Range (µm) | 2.4 to 0.8 | | | | | | | 2.4 to 0.8 | | | | | |
| Pixel size (Å) | 1.08 | | | | | | | 1.08 | | | | | |
| Reconstruction software | cryoSPARC | | | | | | | cryoSPARC | | | | | |
| Micrographs used for data processing | 5,738 | | | | | | | 8,079 | | | | | |
| Symmetry imposed | C1 | | | | | | | C1 | | | | | |
| Initial particle images (no.) | 2,382,280 | | | | | | | 3,833,539 | | | | | |
| Final particle images (no.) | 1,021,093 | 562,605 | 265,043 | 193,445 | 38,065 | 21,565 | 13,827 | 1,262,769 | 554,323 | 362,191 | 346,255 | 158,322 | 90,219 |
| Map resolution (Å) | 3.0 | 3.0 | 3.2 | 3.4 | 3.8 | 4.0 | 4.2 | 3.1 | 3.1 | 3.3 | 3.3 | 3.4 | 3.5 |
| FSC threshold | | | | | | | | | | | | | |
| **Model composition** | | | | | | | | | | | | | |
| Nonhydrogen atoms | 24806 | 24849 | 24835 | 24749 | | | | 24964 | 25051 | 24964 | 24796 | | |
| Protein residues | 3097 | 3105 | 3105 | 3092 | | | | 3123 | 3132 | 3123 | 3123 | | |
| **R.M.S. deviations** | | | | | | | | | | | | | |
| Bond lengths (Å) | 0.004 | 0.004 | 0.007 | 0.004 | | | | 0.006 | 0.004 | 0.005 | 0.004 | | |
| Bond angles (°) | 0.722 | 0.747 | 0.911 | 0.751 | | | | 0.784 | 0.784 | 0.847 | 0.773 | | |
| **Validation** | | | | | | | | | | | | | |
| MolProbity score | 1.28 | 1.39 | 1.48 | 1.45 | | | | 1.37 | 1.46 | 1.62 | 1.48 | | |
| Clashscore | 2.34 | 2.62 | 3.26 | 2.77 | | | | 2.98 | 3.19 | 3.46 | 3 | | |
| Rotomer outliers (%) | 0 | 0 | 0 | 0 | | | | 0 | 0 | 0 | 0 | | |
| **Ramachandran plot** | | | | | | | | | | | | | |
| Favored regions (%) | 96.1 | 95.04 | 94.71 | 94.46 | | | | 95.88 | 94.92 | 92.21 | 94.16 | | |
| Ourliers (%) | 0 | 0 | 0 | 0.03 | | | | 0 | 0.1 | 0.03 | 0 | | |

| | S-GSAS/EG.5.1 | | | | | | | |
|---|---|---|---|---|---|---|---|---|
| | $E1^{1}$ | $E1^{2}$ | $E1^{3}$ | $E1^{4}$ | $E1^{5}$ | $E1^{6}$ | $E1^{7}$ | $E1^{8}$ |
| **PDB ID** | **9AYW** | **9AYX** | **9AYY** | | | | | |
| **EMDB ID** | **44000** | **44001** | **44002** | **44003** | **44004** | **44005** | **44006** | **44007** |
| **Data Collection and processing** | | | | | | | | |
| Microscope | FEI Titan Krios | | | | | | | |
| Detector | Gatan K3 | | | | | | | |
| Magnification | 81000 | | | | | | | |
| Voltage (kV) | 300 | | | | | | | |
| Electron exposure (e-/Å^2) | 57.9 | | | | | | | |
| Defocus Range (µm) | 2.4 to 0.8 | | | | | | | |
| Pixel size (Å) | 1.08 | | | | | | | |
| Reconstruction software | cryoSPARC | | | | | | | |
| Micrographs used for data processing | 11,757 | | | | | | | |
| Symmetry imposed | C1 | | | | | | | |
| Initial particle images (no.) | 4,961,369 | | | | | | | |
| Final particle images (no.) | 289,881 | 258,265 | 154,338 | 187,626 | 126,058 | 48,147 | 39,030 | 23,998 |
| Map resolution (Å) | 3.0 | 3.1 | 3.1 | 3.1 | 3.2 | 3.4 | 3.6 | 4.4 |
| FSC threshold | 0.143 | | | | | | | |
| **Model composition** | | | | | | | | |
| Nonhydrogen atoms | 24813 | 24813 | 24813 | | | | | |
| Protein residues | 3123 | 3123 | 3123 | | | | | |
| **R.M.S. deviations** | | | | | | | | |
| Bond lengths (Å) | 0.004 | 0.004 | 0.004 | | | | | |
| Bond angles (°) | 0.765 | 0.689 | 0.711 | | | | | |
| **Validation** | | | | | | | | |
| MolProbity score | 1.35 | 1.37 | 1.44 | | | | | |
| Clashscore | 2.83 | 3.1 | 3.69 | | | | | |
| Rotomer outliers (%) | 0 | 0 | 0 | | | | | |
| **Ramachandran plot** | | | | | | | | |
| Favored regions (%) | 96.01 | 96.01 | 95.94 | | | | | |
| Ourliers (%) | 0 | 0 | 0 | | | | | |
